# Paneth cell-derived Notch ligands DLL1 and DLL4 are crucial for intestinal crypt base stem cells

**DOI:** 10.64898/2026.09.14.751473

**Authors:** Michaela Quintero, Theresa M. Keeley, Justin Colacino, Nan Gao, Peter J. Dempsey, Linda C. Samuelson

## Abstract

Notch signaling is essential for maintaining intestinal stem cell activity and directing epithelial cell fate. Paneth cells have been proposed to be niche cells, providing Notch signal to neighboring stem cells through expression of the key Notch ligands DLL1 and DLL4. However, intestinal stem cells persist in the absence of Paneth cells. Here, we used genetic mouse models to clarify the role of Paneth cell-derived Notch ligands for stem cell function. Paneth cell-specific deletion of *Dll1* and *Dll4* resulted in loss of crypt base stem cells, with no effect on overall crypt cell proliferation. Organoid growth was reduced in knockout mice, suggesting reduced stem cell function. Upon irradiation injury, stem cell return and crypt regeneration were significantly impaired in the Notch ligand-deleted mice. These findings suggest that Paneth cell-derived Notch ligands DLL1 and DLL4 are crucial for crypt base stem cell maintenance and for crypt regeneration after injury.

## Introduction

Intestinal stem cells preserve the intestinal barrier by continuously renewing the epithelial layer. The *Lgr5*-expressing crypt base columnar cell (CBC), aptly named for its residence in the base of the crypts of Lieberkühn, comprises the principal stem cell population for the mammalian small intestine at homeostasis ^1^. CBCs regularly self-renew and generate highly proliferative, transit-amplifying progenitor cells that dwell within the upper crypt and differentiate into diverse cell types, with epithelial renewal occurring every few days. In addition, due to the remarkably high level of cellular plasticity within the intestinal crypt, many other cell types (including both progenitors and differentiated cells) can act as facultative stem cells when CBC stem cells are lost or impaired ^2–7^. This process involves cellular reprogramming into a stem-like proliferative state to regenerate the tissue, which is accompanied by a specific “fetal-like reversion” gene expression signature (*Ly6a, Ly6d, Ly6e, Clu, Anaxa1, Anexa5, Anexa6*) ^8–10^. The reprogrammed cells occupy open stem cell niche positions at the crypt base to become CBCs, with upregulation of CBC markers, such as *Lgr5* ^2, 6, 11^. Since CBCs are sensitive to many types of injury, including irradiation and inflammation, facultative stem cell populations are thought to be responsible for intestinal repair after stem cell loss to return the tissue to homeostasis ^11^.

Several key niche factors have been identified to maintain CBC function. Of these, Notch signaling has been demonstrated to be essential for directing CBC self-renewal and for lineage commitment to absorptive enterocytes ^12–14^. Acute Notch inhibition pauses stem cell activity, while continued Notch inhibition over several days depletes the stem cell pool through preferential differentiation into secretory cells, including goblet, endocrine, and Paneth cells, while eventually exhausting enterocytes ^12, 15^. Hence, prolonged Notch inhibition ultimately leads to animal death due to the inability to replenish the epithelium ^16^. Notch signaling is also crucial for crypt repair after irradiation-induced CBC injury ^17^.

Notably, while other key stem cell niche factors, such as WNT and growth factors, are derived from multiple cell sources in and around the crypts, Notch signaling requires cell-cell contact for activation ^18, 19^. Thus, Paneth cells are assumed to be the source of the Notch niche signal as they neighbor each CBC and express the key Notch ligands delta like ligand 1(DLL1) and delta like ligand 4 (DLL4), whereas CBCs express the NOTCH1 receptor ^15, 20, 21^. However, functional studies confirming Paneth cell to stem cell Notch signaling are lacking. Moreover, in opposition to this assumption, CBCs remain intact at the crypt base after complete Paneth cell ablation, implying that Paneth cell-derived Notch ligands are dispensable for intestinal stem cell maintenance ^22–25^. It remains unknown how important the Paneth cell-derived Notch signal is for CBC function and how stem cells might compensate for the loss of that signal.

In this study, we aimed to define the Paneth cell-specific role in the Notch niche and thus, generated a Paneth cell-specific Notch ligand ablation mouse model. Surprisingly, after deletion of DLL1 and DLL4 from Paneth cells, CBCs were lost while proliferating cells in the upper crypt remained intact. Additionally, an increased number of Paneth cells filled the crypt base, perhaps implying a drive to generate new Paneth cells to replace the loss of Notch niche cells. We also observed reduced organoid-forming efficiency *ex vivo* and impaired regeneration after irradiation-induced injury *in vivo* in Paneth cell-specific Notch ligand ablated mice. Altogether, our results demonstrate a requirement for Paneth cell Notch ligands to maintain intestinal stem cell function and to regenerate crypts after injury.

## Results

### CBC loss after Paneth cell-specific Notch ligand ablation

To define Paneth cell Notch niche cell function in the intestinal epithelium, we generated mice that deleted *Dll1* and *Dll4* from Paneth cells by crossing *Dll1^fl/fl^*; *Dll4^fl/fl^* mice to a constitutive Paneth cell Cre driver strain (*Defa4^IRES-Cre^; ROSA26-LSL-tdTomato*), which we termed DKO mice. Either Cre-driver mice without floxed alleles or double-floxed mice without the Cre allele were used as controls. We first examined the effect of Paneth cell-specific Notch ligand deletion on intestinal crypt architecture, epithelial cell proliferation, and crypt base columnar (CBC) stem cell maintenance compared to control mice. DKO mice were viable with apparently normal gross intestinal morphology shown by H&E staining (Figure 1A, Figure S1A, B), and normal numbers of proliferating cells, as shown by morphometric counting of EdU-positive cells (Figure 1B). These findings suggested normal intestinal stem cell activity at homeostasis. However, staining for the CBC marker olfactomedin 4 (OLFM4) revealed a loss of cells expressing both mRNA and protein from the crypt base in DKO mice (Figure 1C, Figure S2A). Since *Olfm4* is a Notch target gene ^12^, we also analyzed other CBC markers unaffiliated with Notch signaling. Solute carrier family 12 member 2 (SLC12A2), which normally marks CBC stem cells and transit-amplifying progenitors, was also lost from the crypt base yet retained in the upper crypt of DKO mice (Figure 1D). Similarly, mRNA expression of the CBC marker and WNT target *Ascl2* and the CBC marker *Hmgcs2,* which shows excellent specificity to CBCs ^26^, showed reduced expression of *Ascl2* at the crypt base and an almost complete loss of *Hmgcs2* (Figure 1E). Additionally, consistent with the histological analysis, mRNA abundance of CBC markers *Olfm4, Notch1, Lgr5,* and *Ascl2* was reduced in DKO intestine (Figure 1F). Further, mRNA expression of the Notch target gene *Hes1* was also significantly reduced, consistent with reduced Notch signaling in the DKO crypts (Figure S2B).

**Figure 1.**
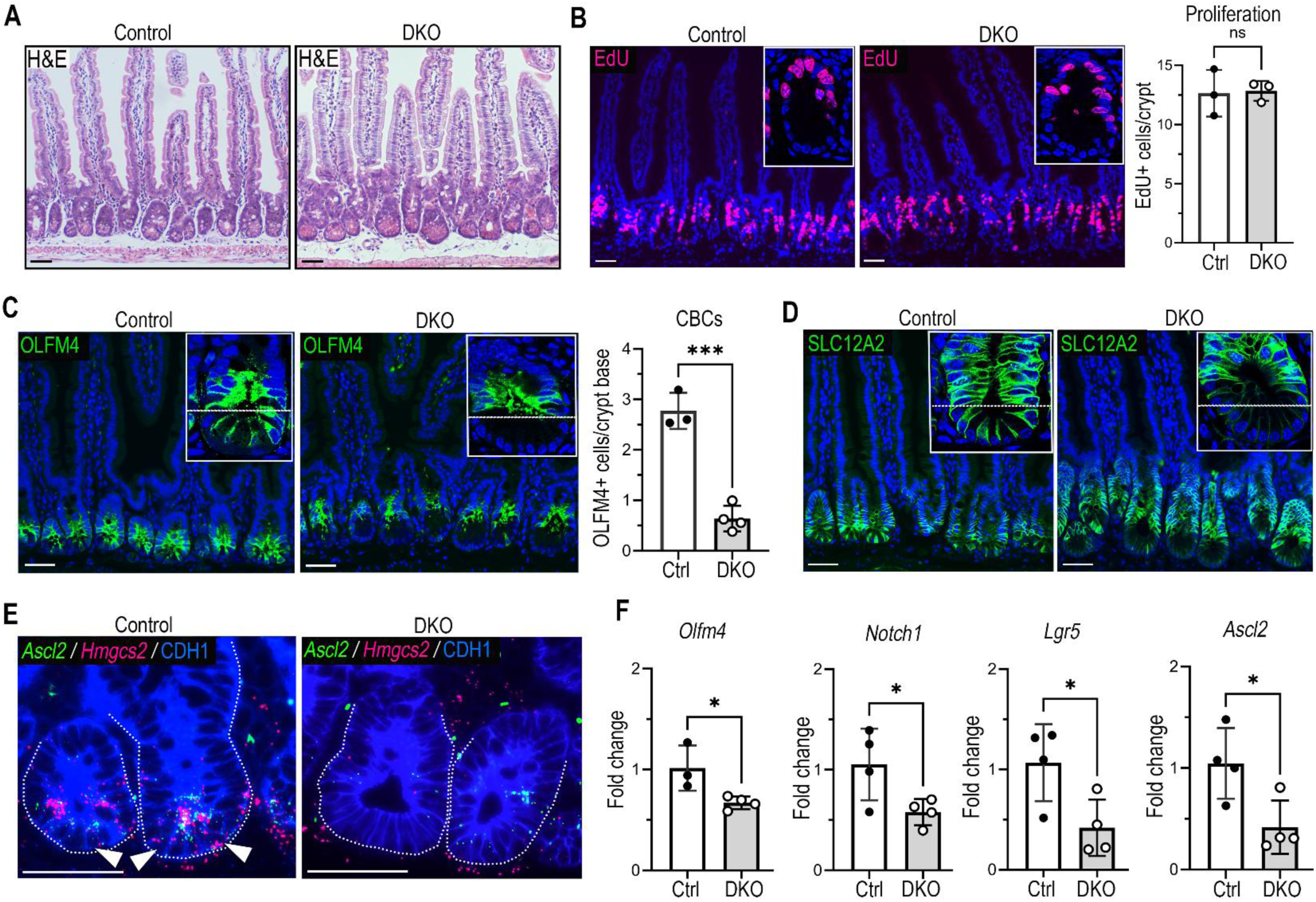
CBCs are lost from the crypt base after Notch ligand depletion from Paneth cells. (A) Hematoxylin and eosin (H&E) staining of the jejunum from control and DKO mice (*Dll1^f/f^; Dll4^f/f^; Defa4^IRES-Cre^; ROSA26-LSL-tdTomato*). Scale bars, 50 μm. (B) EdU (magenta) staining of jejunal sections with nuclear DAPI staining (blue). Mice were injected with EdU 1.5 hours prior to tissue collection. Insets are separate confocal images of crypts. EdU^+^ cells were counted for 30-60 well-oriented jejunal crypts per mouse. Quantitative data are presented as mean ± SD (control (Ctrl) versus DKO by unpaired two-tailed Welch’s t test; ns, not significant; n = 3 mice/group). Scale bars, 50 μm. (C) Immunofluorescence (IF) staining of crypt base columnar (CBC) stem cell marker and Notch target OLFM4 (green) with DAPI (blue). Insets are confocal images of crypts. OLFM4+ cells below the +4 region (marked by dotted white line) were counted for all well-oriented crypts within a 4 cm strip of jejunum per mouse (60-130 crypts). Quantitative data are presented as mean ± SD (***p < 0.0001, Ctrl versus DKO by unpaired two-tailed Student’s t test; n = 3-4 mice/group). Scale bars, 50 μm. (D) IF staining of CBC and crypt cell marker SLC12A2 (green) with DAPI (blue). Insets are confocal images of crypts with dotted white line marking the base. Scale bars, 50 μm. (E) In situ hybridization for CBC markers *Ascl2* (green) and *Hmgcs2* (magenta) with CDH1 (E-cadherin) IF staining (blue). Arrowheads point to crypt base cells expressing both *Ascl2* and *Hmgcs2*. Scale bars, 50 μm. (F) mRNA abundance of CBC markers was measured by qRT-PCR analysis of jejunal crypt RNA. Quantitative data are presented as mean fold-change ± SD (Ctrl versus DKO by unpaired two-tailed Student’s t-test or Welch’s t-test when variance between DKO and controls differed; *p < 0.05; n = 3-4 mice/group).

Notably, CBC loss from the crypt base was observed throughout the intestine, with penetrance increasing from the proximal to distal small intestine, which reflected the expression pattern of the Cre driver as demonstrated by the prevalence of tdTomato-labeled Paneth cells in the DKO mouse model (Figure S1C). Large intestinal CBCs remained unaffected, as expected from the restricted expression domain of *Defa4^IRES-Cre^* to the small intestine (Figure S1D). The normal proliferation and intestinal morphology in DKO mice suggest that facultative stem cell populations in the upper crypt compensate for CBC absence to maintain the intestinal epithelium.

### Inducible ablation of Dll1 and Dll4 in Paneth cells triggers crypt cell death

To study the acute stem cell changes underlying crypt remodeling after Paneth cell Notch ligand deletion, we generated an inducible model by crossing *Dll1^fl/fl^*; *Dll4^fl/fl^* mice to a tamoxifen-inducible Paneth cell Cre driver strain (*Lyz1^3’UTR-IRES-CreERT2^*), which we termed IDKO mice. Ligand deletion was induced in Paneth cells of adult mice by tamoxifen (TAM) treatment for 5 days, with intestines harvested 2 days later (Figure 2A). TAM-treated *Lyz1^3’UTR-IRES-CreERT2^*mice served as controls. Like the DKO constitutive model, acute deletion of DLL1 and DLL4 in Paneth cells reduced the number of OLFM4-expressing and SLC12A2-expressing cells at the crypt base (Figure 2B, C). Strikingly, H&E staining revealed numerous crypts with delaminating cells in IDKO mice, suggesting an upsurge in cell death after acute Notch ligand deletion from Paneth cells (Figure 2D, arrowheads). Accordingly, cleaved caspase 3 (CC3) staining showed an increased number of apoptotic cells at the crypt base in IDKO mice compared to controls (Figure 2E). Co-staining for CC3 and the Paneth cell marker matrix metalloprotease 7 (MMP7) showed that apoptotic cells included both Paneth cells and neighboring crypt cells (Figure 2E). To capture an earlier stage, we treated some IDKO mice with a single dose of TAM and harvested crypts three days later, with similar results (Figure S3A, B). These findings align with our previous study that defined the consequences of acute Notch inhibition using a pharmacological approach, which induced rapid Paneth cell death while inhibiting stem cell activity ^15^. Our analysis of the IDKO model showed that cell death is associated with the crypt cell remodeling and CBC stem cell loss induced by ablating Notch signals from Paneth cells, suggesting that Paneth cell niche support is important for CBC survival.

**Figure 2.**
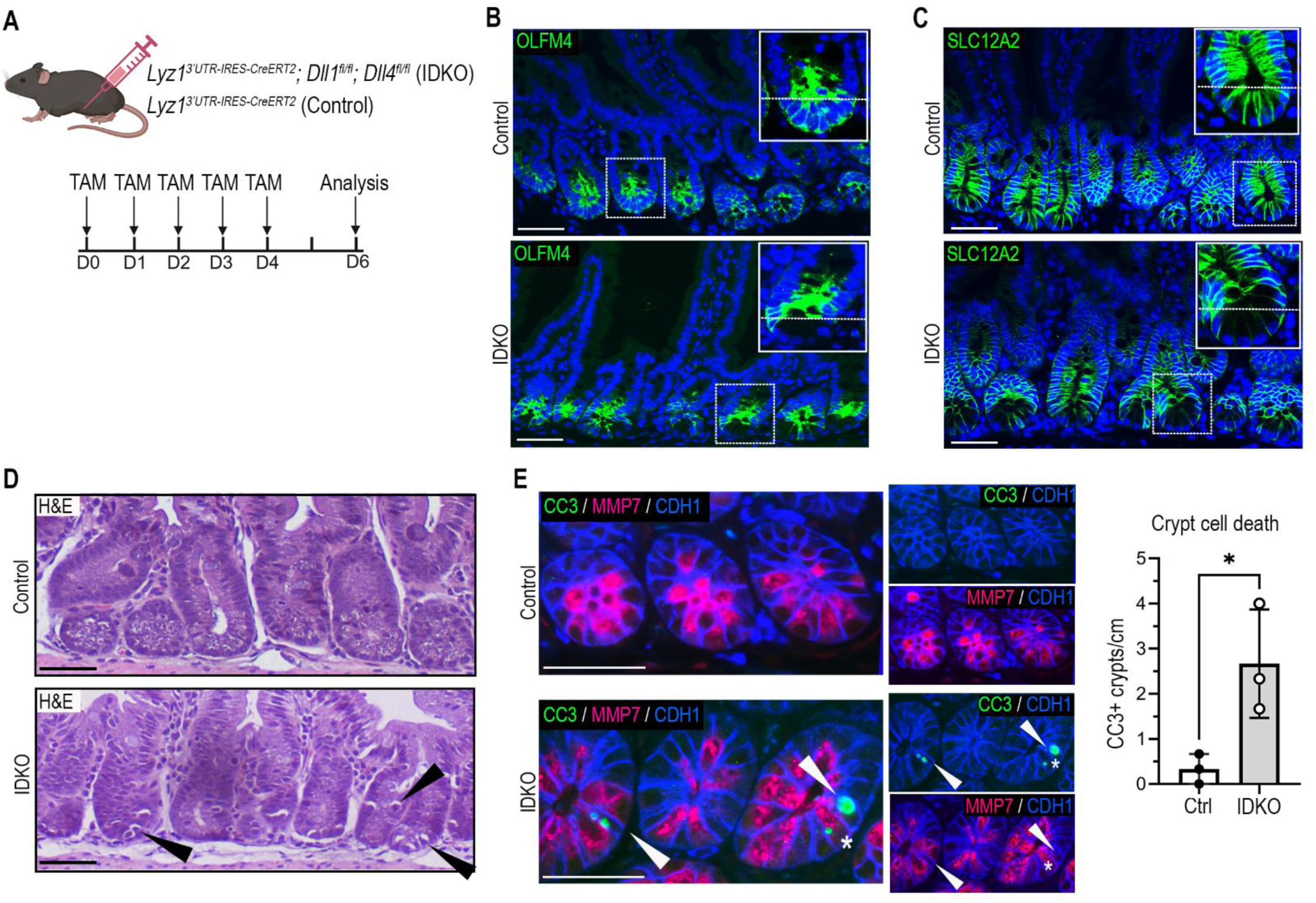
Acute Notch ligand depletion from Paneth cells induces crypt cell death. (A) Schematic of experimental design. Adult IDKO mice (*Lyz1^3’UTR-IRES-CreERT2^*; *Dll1^fl/fl^*; *Dll4^fl/fl^*) or control mice (*Lyz1^3’UTR-IRES-CreERT2^)* were treated with tamoxifen (TAM) for 5 days and intestines were harvested 2 days later for analysis. (B) IF staining for OLFM4 (green) with DAPI (blue) on jejunal sections. Insets show marked crypts at higher power. Scale bars, 50 μm. (C) IF staining for SLC12A2 (green) with DAPI (blue) on jejunal sections. Insets show marked crypts at higher power. Scale bars, 50 μm. (D) H&E staining jejunal tissue sections. Arrowheads point to examples of delaminating cells. Scale bars, 50 μm. (E) IF staining for cleaved caspase 3 (CC3; green), Paneth cell marker MMP7 (magenta), and CDH1 (blue). Arrowheads point to MMP7-negative/CC3-positive cells, while asterisks show examples of MMP7- positive/CC3-positive cells. Quantitative data are presented as mean ± SD (Ctrl versus IDKO by unpaired two-tailed Welch’s t test; *p < 0.05; n = 3 mice/group). Crypts containing CC3^+^ cells were counted from well-oriented crypts within a 3 cm strip of jejunum per mouse measured in ImageJ. Scale bars, 50 μm.

### Paneth cell expansion after Paneth cell-specific Notch ligand ablation

A focus on Paneth cell changes in the constitutive model revealed a marked expansion of Notch ligand-depleted cells at the DKO crypt base, with increased numbers of cells expressing the Paneth cell marker lysozyme (LYZ) compared to control (Figure 3A). There were also numerous mislocalized tdTomato-labeled “Paneth cells” within the villi of DKO mice (Figure 3B, arrowheads), which co-expressed both Paneth cell (LYZ) and goblet cell (mucin-2, MUC2) markers (Figure S4A, B, arrowheads). This co-expression phenotype is a feature of intermediate cells, which have been described as Paneth cell precursors ^27, 28^. Accordingly, co-staining for another Paneth cell marker, MMP7, with MUC2 confirmed that the mislocalized “Paneth cells” on the intestinal villi exhibited intermediate cell features (Figure 3C, arrowheads). The MMP7-positive cells on the villus expressed both markers, while cryptal MMP7-positive cells lacked MUC2, as expected for mature Paneth cells (Figure 3C). The substantial increase in Paneth and intermediate cells suggests that DKO mice may possess a drive to generate new Paneth cells to replace the dysfunctional ones lacking Notch ligands.

**Figure 3.**
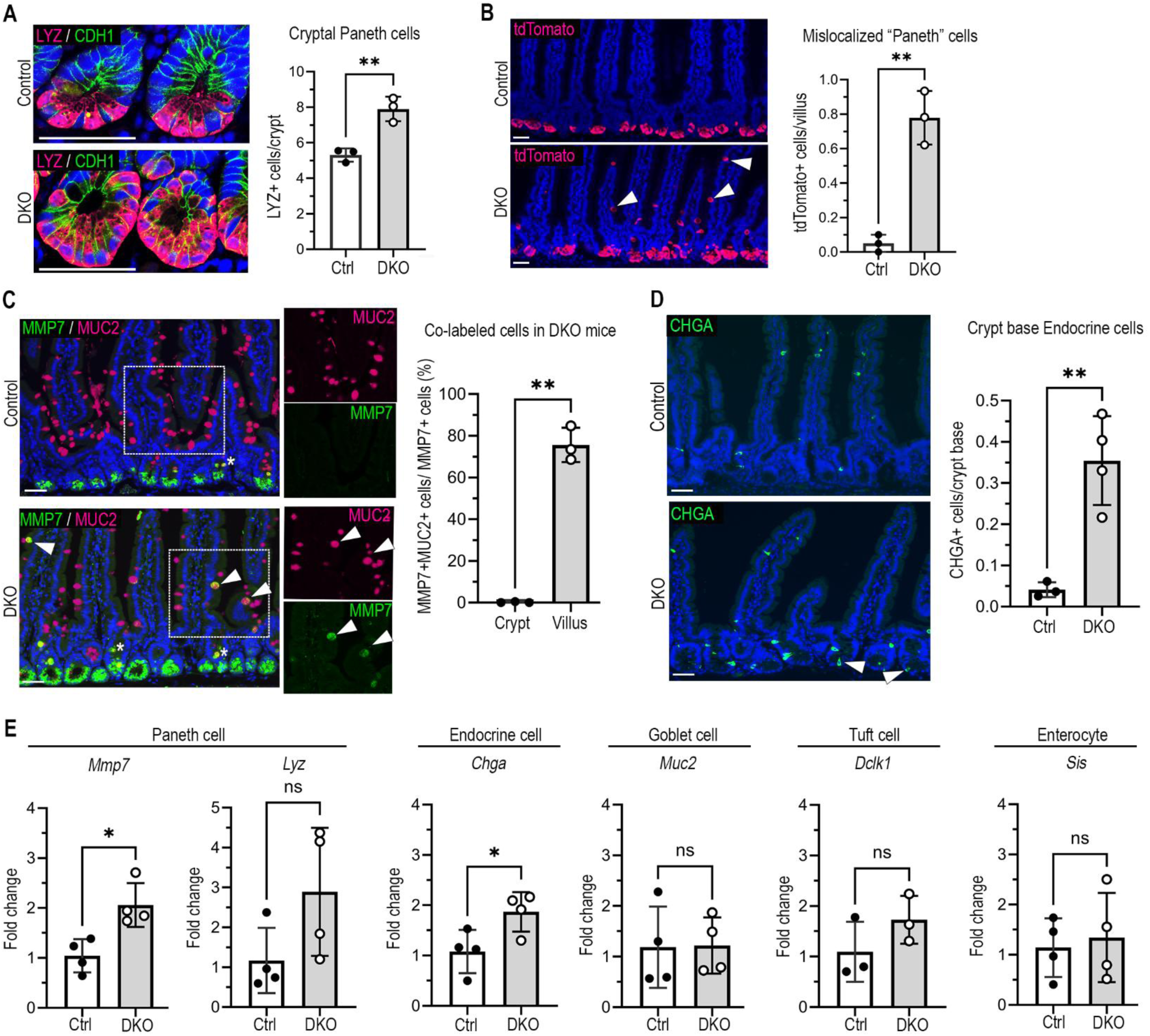
Notch ligand depletion from Paneth cells drives Paneth cell expansion. (A) Confocal imaging of jejunal crypts after IF staining for lysozyme (LYZ, magenta) and CDH1 (green), with DAPI (blue) on control and DKO tissue sections. LYZ^+^ cells were counted from 60-200 well-oriented jejunal crypts per mouse. Quantitative data are presented as mean ± SD (Ctrl versus DKO by unpaired two-tailed Student’s t-test; **p < 0.001; n = 3 mice/group). Scale bars, 50 μm. (B) IF staining for tdTomato (magenta) with DAPI (blue) in jejunal tissue sections. tdTomato^+^ cells were counted from 100-200 well-oriented villi per mouse. Quantitative data are presented as mean ± SD (Ctrl versus DKO by unpaired two-tailed Welch’s t test; **p < 0.001; n = 3 mice/group). Scale bars, 50 μm. (C) IF staining for Paneth cell marker MMP7 (green) and goblet cell marker MUC2 (magenta) with DAPI (blue). Arrowheads denote co-labeled cells on the villus. Asterisks denote co-labeled cells in the upper crypt where intermediate cells are located. Outsets show green or magenta channels from the boxed region. MMP7^+^ or MMP7^+^/MUC2^+^ co-labeled cells were counted in 100-200 well-oriented crypts and villi per mouse. For crypt counts, only co-labeled cells within the base of the crypt were counted. Quantitative data are presented as mean ± SD (% of co-labeled cells in the crypt versus villus in DKO mice by unpaired two-tailed Welch’s t test; *p < 0.01; n = 3 mice/group). Scale bars, 50 μm. (D) IF staining for endocrine cell marker chromogranin A (CHGA, green) in jejunal sections with DAPI (blue). Insets are enlarged images of selected regions. Arrowheads point to CHGA^+^ cells within the crypt base. Quantitative data are presented as mean ± SD (Ctrl versus DKO by unpaired two-tailed Welch’s t test; *p < 0.01; n = 3 mice/group). CHGA^+^ cells below the +4 region of the crypt were counted for 100-140 crypts well- oriented jejunal crypts per mouse. Scale bar, 50 μm. (E) Quantification of mRNA abundance of differentiated cell markers in jejunal crypt RNA by RT-qPCR. Quantitative data are presented as mean ± SD (^∗∗^Ctrl versus DKO by unpaired two-tailed Welch’s t-test; *p < 0.05; n = 4 mice/group). Scale bar, 50 μm.

To assess whether there were changes to the differentiation of other intestinal epithelial cell types we analyzed DKO and control mice for cell-specific markers by immunofluorescence and reverse transcriptase quantitative polymerase chain reaction (RT-qPCR). Interestingly, staining for the pan-endocrine marker chromogranin A (CHGA) showed increased numbers of endocrine cells present at the crypt base (Figure 3D). However, CHGA-expressing crypt-base cells were sparse, averaging less than one cell per crypt analyzed. Consistent with the histological analysis, RT-qPCR showed increased expression of Paneth cell (*Mmp7)* and endocrine cell (*Chga*) markers in DKO crypts (Figure 3E). In contrast, overall numbers of goblet cells, assessed by MUC2 or AB/PAS staining, and tuft cells, assessed by DCLK1 staining, remained unaltered (Figure S4D-F). Likewise, mRNA abundance of goblet cell (*Muc2*), tuft cell (*Dclk1*), and enterocyte (*Sis*) markers was normal in DKO crypts (Figure 3E). Thus, other than CBCs and Paneth cells, the overall cell census is largely normal in the DKO intestine.

### Paneth cells transcriptionally remodel after Notch ligand deletion

We examined whether DKO Paneth cells altered expression of other stem cell niche factors to compensate for loss of Notch ligands by performing bulk RNA-sequencing (RNA-seq) on FACS-isolated EPCAM/tdTomato co-labeled Paneth cells from DKO and control mice (Figure 4A, Figure S5A). Villi were removed prior to crypt isolation to enrich for cryptal Paneth cells to minimize the contribution of DKO tdTomato-labeled cells from the villus. RNA-seq analysis of sorted Paneth cells showed separation between DKO and control samples by multidimensional scaling (MDS) (Figure S5B). Common Paneth cell mRNAs were highly expressed and unchanged in DKO compared to control, including *Lyz1, Mmp7, Mtpx2,* and 25 defensin genes, confirming the Paneth cell identity of sorted cells with no apparent changes in antimicrobial gene expression (Figure 4B, Table S1). Importantly, DKO Paneth cells also maintained normal expression levels of non-Notch niche factor mRNAs, including canonical WNT ligand *Wnt3* and the growth factors *Tgfa*, *Egf*, *Ereg*, and *Areg* (Figure 4B-C, Table S1).

**Figure 4.**
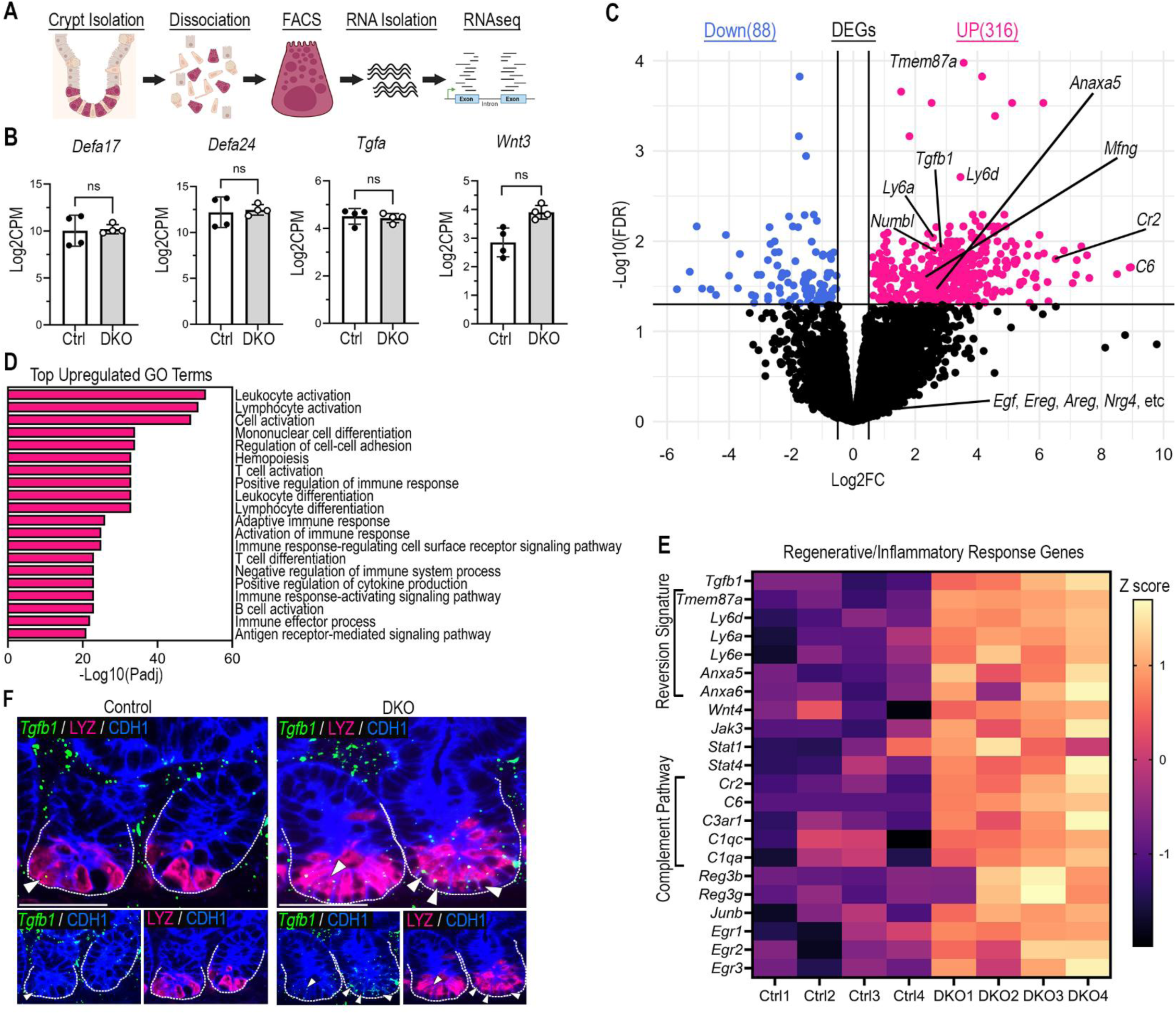
Transcriptional remodeling of Paneth cells in DKO mice. (A) Paneth cells from Ctrl and DKO mice were isolated by FACS by gating on Epcam and tdTomato, followed by transcriptomics analysis by bulk RNA-sequencing (RNAseq). See Figure S5A for FACS plots. (B) Normalized counts per million (log2CPM) of highly expressed and unchanged genes in control (Ctrl) and DKO mice. Quantitative data are presented as mean ± SD; significance was determined by FDR-adjusted p-value; ns, not significant. (C) Volcano plot of RNAseq data showing differentially expressed genes (DEGs) in DKO versus Ctrl Paneth cells, with 316 significantly upregulated genes (magenta) and 88 downregulated genes (blue). Selected upregulated genes mentioned in the text are labeled. The complete list of DEGs are listed in Table S2. (D) Selected upregulated GO terms from analysis of the top 100 upregulated DEGs ranked by F score. The complete list of GO terms are listed in Table S3. (E) Heatmap of select upregulated DEGs involved in tissue regeneration. The intestinal fetal reversion signature and complement pathway genes have been described to be associated with regenerative potential (see text). (F) In situ hybridization for the growth factor mRNA *Tgfb1* by RNAscope (green) with IF co-staining for LYZ (magenta) and CDH1 (blue), showing increased *Tgfb1* expression in DKO Paneth cells. Outsets show green (*Tgfb1)* or magenta (LYZ) channels with CDHI. Arrowheads indicate *Tgfb1*^+^ and LYZ^+^ co-positive cells. Scale bars, 50 μm.

Further analysis revealed several differentially expressed genes (DEGs), with 316 upregulated and 88 downregulated genes in DKO Paneth cells compared to control Paneth cells (Figure 4C; Table S2). The Notch modulators, manic fringe (*Mfng*) and numb-like endocytic adaptor protein *(Numbl*), were among the upregulated DEGs, suggesting an attempt to regulate Notch components after DLL1 and DLL4 ligand loss (Figure 4C). Among other upregulated DEGs were transcription factors involved in Paneth cell lineage commitment and maturation, *Gfi1* and *Bhlha15* (*Mist1*) ^27, 29^ (Figure S5C). Gene ontology (GO) term analysis revealed that most of the upregulated genes were associated with the regulation of inflammation and the adaptive immune response (Figure 4D, Table S3); whereas downregulated genes displayed more modest changes with unclear function (Figure S5D). Deeper analysis revealed that many of the most highly upregulated DEGs were associated with tissue repair and the protective side of inflammation. For instance, *Tgfb1,* which acts both as a stem cell niche factor and a key regulator of regeneration ^10^, was upregulated roughly 7-fold in DKO Paneth cells (Figure 4C, E, Table S2). RNAscope staining for *Tgfb1* mRNA confirmed increased expression in DKO Paneth cells (Figure 4F). Furthermore, DKO Paneth cells upregulated expression of regenerative genes sometimes described as a “fetal reversion” signature that act downstream of TGFB1 and/or YAP/TAZ and are thought to underlie the activation of facultative stem cells during crypt regeneration ^8–10, 30^, including *Ly6a, Ly6d*, *Ly6e*, *Anxa5*, *Anxa6*, and *Tmem78a,* (Figure 4C, E).

Interestingly, components of the complement system, including *C6* and *Cr2*, were among the most highly upregulated genes in DKO Paneth cells (Figure 4E, Table S2). Notably, complement system upregulation in niche cells has been shown to promote stem cell function in the intestine and other tissues ^31–33^. Accordingly, *Wnt4*, a non-canonical WNT ligand associated with the regenerative response ^34, 35^ was also upregulated (Figure 4E, Table S2). Additionally, other inflammation-associated upregulated DEGs, such as *Jak1*, *Stat1*, *Reg3b, Reg3g, Junb,* and *Egr1*, have been associated with the support of epithelial regeneration and/or wound healing ^36–39^ (Figure 4E, Table S2). Although many inflammation-related genes were differentially expressed in DKO Paneth cells, RT-qPCR analysis of crypt RNAs showed no differences in expression of the damage-associated inflammatory markers *Infg*, *Il1b*, and *Tnfa,* suggesting that DKO crypts did not adopt an inflammatory tone (Figure S5E). We also assessed inflammatory cell infiltration by CD45 staining and observed no apparent differences in immune cell numbers or epithelial infiltration between DKO and control (Figure S5F). Altogether, this analysis showed that DKO Paneth cells retained expression of antimicrobial peptides and homeostatic stem cell support genes after Notch ligand deletion, while increasing expression of genes associated with stem cell regeneration after injury.

### Reduced organoid-forming efficiency in DKO mice

Because genes involved in supporting regeneration were upregulated in DKO Paneth cells, we evaluated stem cell function by comparing control and DKO organoid establishment efficiency of single crypt cells plated into high-WNT culture media (50% L-WRN). Notably, DKO crypt cells exhibited decreased organoid-forming efficiency compared to control cells, indicating an epithelial cell-intrinsic decline in stem cell activity when Notch ligands are deleted from Paneth cells (Figure 5A). In addition, DKO organoids appeared smaller with slightly irregular morphology compared to control organoids, which take on a regular cystic morphology in high-WNT culture conditions. We next tested the ability to form mature, budding enteroids by plating intestinal crypts and growing in ENR differentiation media. Interestingly, DKO enteroids failed to form buds, while control organoids grew into large complex structures with many buds (Figure 5B). Furthermore, tdTomato+ Paneth cells within DKO organoids clustered together within the main enteroid body, while tdTomato+ cells were dispersed within the buds of control organoids consistent with normal crypt cellular architecture. These results demonstrate a reduced capacity of DKO crypt cells to establish enteroids and suggest a lack of ability for DKO enteroids to form crypt-like structures.

**Figure 5.**
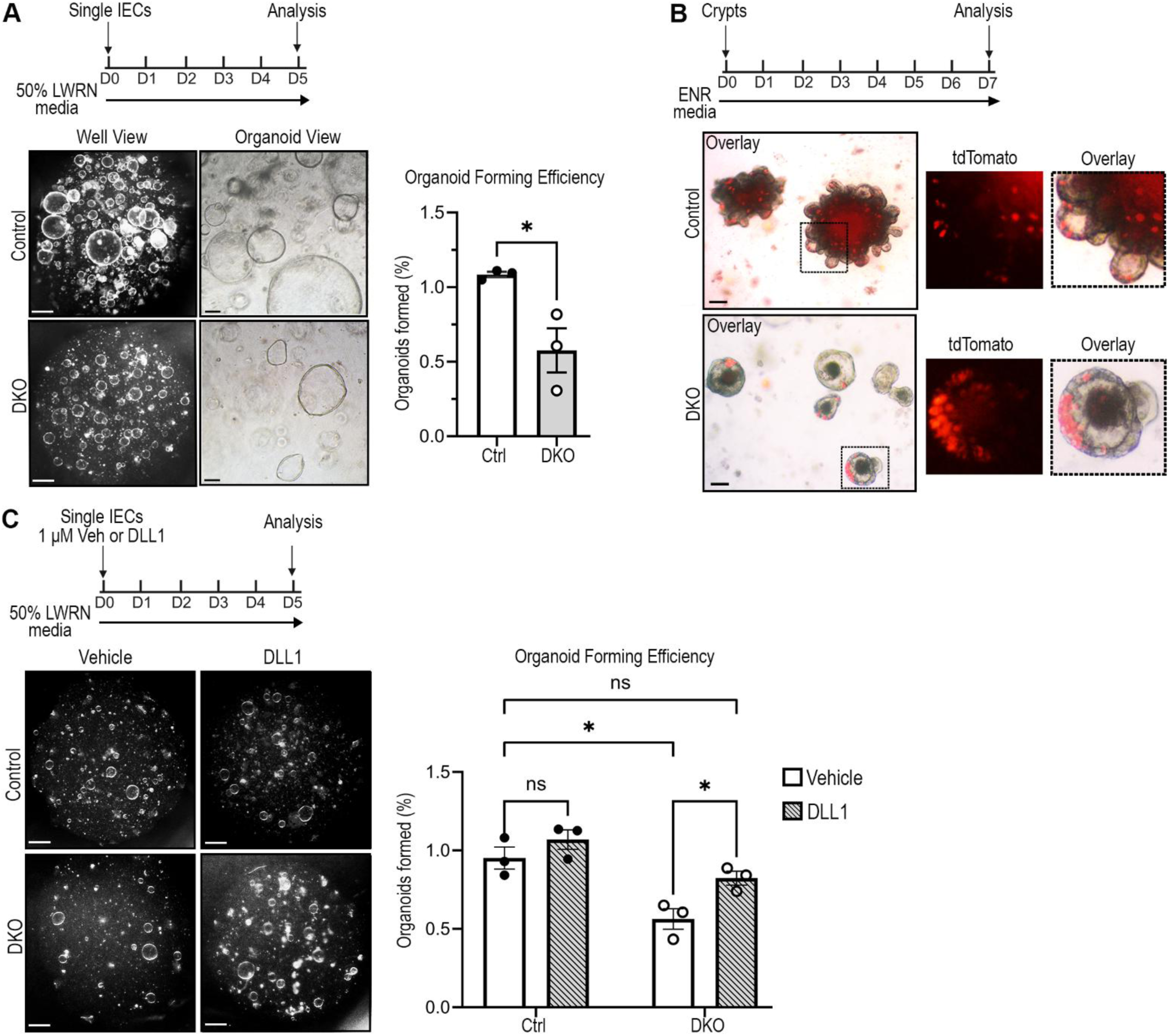
Reduced organoid-forming efficiency and organoid crypt budding in DKO mice. (A) Organoid forming efficiency was measured in control and DKO mice. Dispersed jejunal crypt cells were plated in Matrigel at 1,200 cells/µL and the number of organoids formed was counted 5 days later after growth in L-WRN growth media. Views of a single well (left, scale bar, 50 mm) and organoids (right, scale bar, 50 μm) are shown. Quantitative data are presented as mean ± SD of the % of the input cells that formed organoids from 8 wells per mouse (Ctrl versus DKO by unpaired two-tailed Welch’s t test; *p < 0.05; n = 3 mice/group). (B) Jejunal crypts were plated into ENR differentiation media for 7 days to assess growth and differentiation of control and DKO organoids. Increased DKO crypt numbers were plated to match organoid densities in control cultures (see Experimental Procedures). Images show organoid structure with Paneth cells identified by tdTomato fluorescence. Outsets are higher power images of boxed regions, showing the tdTomato channel and the overlay with phase contrast. Scale bars, 50 μm. (C) Organoid forming efficiency was measured after plating single crypt cells seeded at 1,000 cells/µL in Matrigel containing Notch ligand DLL1 (1 μM) or vehicle followed by growth in L-WRN media. Single well views from control and DKO organoid cultures are shown. Scale bar, 50mm. The number of organoids formed is presented as mean ± SD of 6 wells per mouse (Ctrl versus DKO and vehicle versus DLL1 in each mouse group were assessed by two-way ANOVA with Bonferroni’s correction for multiple tests;*p <0.05; n = 3 mice/group).

We next tested whether the decline in organoid formation was a direct consequence of the loss of Notch signals to the stem cells. Organoid-forming capacity was measured with and without the addition of recombinant Notch ligand DLL1 to the Matrigel before plating single crypt cells. Importantly, the addition of DLL1 to the Matrigel significantly increased the number of organoids generated from DKO crypt cells to a rate comparable to controls (Figure 5C). In contrast, the addition of DLL1 did not significantly alter the organoid-forming ability of controls. This result demonstrates that impaired stem cell function in DKO mice can be rescued by the addition of DLL1, implying a direct effect of the Paneth cell-derived Notch ligands on stem cell activity.

### Impaired crypt regeneration in DKO mice

Since Notch signaling is essential for crypt regeneration after irradiation injury ^17^, we tested whether DKO mice properly regenerate crypts after 12 Gy of whole-body irradiation. DKO and control intestines were analyzed histologically during the regenerative phase of the injury response 3 days after irradiation ^40^ (Figure 6A). As expected, control mice displayed elongated, highly proliferative crypts with an expansion of cells co-labeled for the stem cell marker SLC12A2 and the proliferation marker EdU (Figure 6B, C). In stark contrast, DKO crypts were shorter with fewer SLC12A2-marked cells and reduced proliferation (Figure 6B, C). Consistently, crypts housing cells expressing the stem cell marker and Notch target gene *Olfm4* were rare in DKO mice after irradiation, whereas the regenerative crypts in control mice contained many *Olfm4*-expressing cells, indicating that Notch signaling was significantly impaired in DKO intestine after injury (Figure 6D). Both control and DKO mice retained tdTomato-labeled Paneth cells, with no signs of lineage tracing at day 3 after irradiation (Figure 6E). Interestingly, we observed increased mislocalized crypt-like structures in DKO mice after irradiation, as shown by H&E and crypt cell marker staining (Figure 6B, C, E, arrows). Because our gene expression profiling showed that DKO Paneth cells expressed an inflammation-associated signature (Figure 4) that might be related to intestinal dysbiosis, we tested the role of the microbiota in the injury response. We treated DKO and control mice with broad-spectrum antibiotics ^41^ for 2 weeks prior to irradiation injury. Analysis of the intestinal regenerative response showed that antibiotic treatment did not prevent the impaired regeneration in DKO mice, with fewer EdU-, SLC12A2-, and OLFM4-expressing cells in regenerating DKO crypts, consistent with our earlier observation (Figure S6 A-D). Our data indicates that Paneth cell Notch ligands are required to effectively regenerate crypt cells after irradiation injury, suggesting that other Notch ligand-expressing cells in the DKO crypt are unable to compensate for the Paneth cell defect for crypt regeneration.

**Figure 6.**
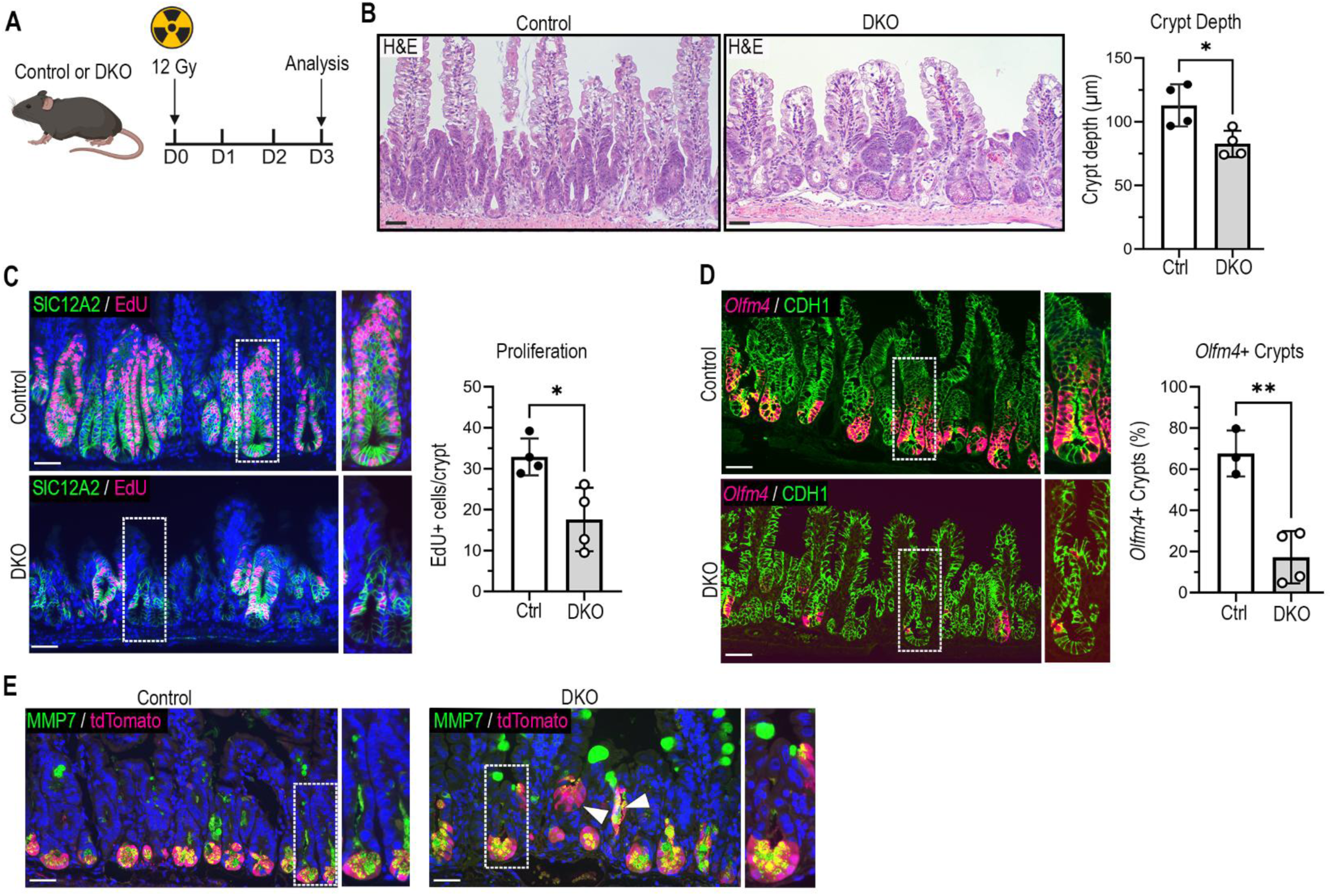
Impaired crypt regeneration after radiation injury in DKO mice. (A) Experimental schematic. Control and DKO mice were treated with whole-body 12 Gy cesium irradiation to induce stem cell injury, and intestines were harvested 3 days later for histological analysis. Mice were treated with EdU 1.5 hours prior to tissue collection. (B) H&E staining of the jejunum from irradiated DKO and control mice. The depth of each crypt within a 4 cm section of jejunum was measured with Image-J software and graphed as mean ± SD (Ctrl versus DKO by unpaired two-tailed Student’s t test; *p < 0.05: n = 4 mice/group). Scale bars, 50 μm. (C) IF staining of jejunum for SLC12A2 (green) and EdU (magenta) with DAPI (blue). Outsets show enlarged images of boxed region. EdU^+^ cells were counted for 30-60 well-oriented jejunal crypts per mouse and presented as mean ± SD (Ctrl versus DKO by unpaired two-tailed Student’s t test; *p < 0.05; n = 4 mice/group). Scale bars, 50 μm. (D) In situ hybridization for CBC marker *Olfm4* (magenta) by RNAscope with IF co-staining for CDH1 (green) in jejunal sections. Outsets show enlarged images of boxed regions. *Olfm4*-positive crypts were counted and graphed as mean ± SD of % of total crypts counted (Ctrl versus DKO by unpaired two-tailed Student’s t test; **p < 0.001; n = 3-4 mice/group). Scale bars, 50 μm. (E) IF staining for MMP7 (green) and tdTomato (magenta) with DAPI (blue). Outsets show enlarged images of boxed regions. Arrowheads point to mislocalized crypts. Scale bars, 50 μm.

## Discussion

Our study revealed the importance of Paneth cell-derived Notch signal for CBC maintenance and intestinal stem cell function. After deletion of the Notch ligands DLL1 and DLL4 from Paneth cells, CBCs were lost from the crypt base, while upper-crypt proliferating cells were apparently responsible for maintaining the epithelium in place of CBCs. We also observed increased numbers of Paneth and intermediate cells in DKO intestine, which may imply a drive to generate new Paneth cells to replace the defective Paneth cells lacking Notch ligands. Our findings highlight the extreme crypt cell plasticity to maintain the intestinal epithelium after stem cell dysfunction caused by disruption of the CBC niche. Furthermore, Paneth cell-specific Notch ligand deletion reduced overall intestinal stem cell function, as demonstrated by decreased organoid-forming efficiency *ex vivo* and impaired intestinal regeneration after irradiation-induced injury *in vivo*. Altogether, our results establish an important role for Paneth cell Notch ligands to maintain intestinal stem cell function and for crypt recovery after injury.

Long recognized as specialized crypt cells important for host defense and innate immunity, the importance of Paneth cells as CBC niche cells has been debated, with some studies suggesting niche function while others showed that intestinal stem cells tolerate complete Paneth cell loss. We found that the loss of DLL1 and DLL4 in Paneth cells depleted CBCs, thus demonstrating that Notch signal from Paneth cells is essential for CBC maintenance. Our finding of CBC loss after Paneth cell Notch-ligand deletion agrees with a study that showed reduced numbers of OLFM4-expressing CBCs after incomplete Paneth cell loss in adult mice ^20^. However, in contrast to CBC loss after Paneth cell Notch ligand deletion, other studies have shown CBC retention after complete Paneth cell or secretory cell ablation, with CBCs instead expanding to fill the crypt base ^22–25^. The conflicting findings regarding CBC loss versus CBC retention imply that separate compensatory mechanisms exist to maintain the intestinal epithelium after total loss of Paneth cells versus targeted loss of Paneth cell Notch ligands. Most importantly, functional support of CBCs by Paneth cells is uncontested, as they markedly improve the capacity of CBCs to form organoids ^20, 42^. Our data reinforces this concept of Paneth cells as key CBC niche cells.

Despite CBC loss, cellular proliferation was normal at homeostasis in DKO intestine, demonstrating cellular remodeling to preserve the epithelium. Cells expressing stem cell markers, including the Notch target OLFM4, persisted within the DKO upper crypt, whereas Notch ligand-depleted Paneth cells expanded in number to fill the base of the crypt, clustering away from the upper-crypt proliferative zone. The pattern of expression of the Notch-target gene *Olfm4* in the DKO intestine indicates that when the normal Notch niche at the crypt base is lost, the crypt remodels to build a new Notch stem cell niche within the upper crypt. This suggests that Notch ligand-expressing secretory progenitors may be sufficient to maintain the Notch niche for alternate OLFM4-expressing intestinal stem cells. This cellular remodeling agrees with previous studies showing that CBCs are dispensable for epithelial renewal due to the inherent plasticity within the intestinal epithelium ^11^. Facultative stem cells have been shown to obtain intestinal stem cell identity after CBC loss or injury ^2–7^, and various alternative homeostatic stem cells, such as *Fgfbp1*-expressing cells, have been proposed in the upper crypt ^43, 44^. The nature of the “stem cells” maintaining the intestinal epithelium within the new Notch niche of the CBC-depleted DKO mice is an interesting future question.

Except for the marked increase in Paneth cells, epithelial cell differentiation was only modestly changed in DKO mice, with apparently normal differentiation of the two most prevalent cell types, goblet cells and enterocytes. However, endocrine cell numbers were increased, with more endocrine cells located with Paneth cells at the crypt base. Increased crypt-base endocrine cells and tuft cells were previously reported as a possible compensatory mechanism to support CBCs after acute Paneth cell depletion ^25^, as these cells, like all secretory cell types, express DLL1 and DLL4 ^15, 25^. Still, endocrine cell increases in our model and after acute Paneth cell depletion were not of sufficient numbers to substitute for Paneth cell Notch niche function. Complete substitution of the Paneth cell Notch niche would require Notch ligand-expressing cells to neighbor each CBC. Moreover, the depletion of CBCs in our model demonstrates that the increased prevalence of crypt base endocrine cells does not prevent CBC loss.

Interestingly, Notch ligand-depleted Paneth cells exhibited unexpected changes. Cryptal Notch ligand-depleted Paneth cells underwent transcriptional remodeling, increasing expression of regenerative genes while retaining mature cell-identity markers, such as antimicrobial genes. Transcriptomic analysis of the Notch ligand-depleted Paneth cells showed that other proposed CBC niche factors, including *Wnt3* and growth factors such as *Tgfa*, *Egf*, *Ereg*, and *Areg*, were expressed at normal levels, thus, reinforcing the specific role of Notch ligands for CBC maintenance. The increased expression of genes encoding signaling factors that have been described to support tissue repair, such as *Tgfb1* and *Wnt4* in DKO mice suggests that Paneth cells potentially upregulate these factors to support crypt remodeling upon CBC loss. WNT4 is a non-canonical WNT ligand that is not normally highly expressed in normal Paneth cells and is potentially involved in crypt fission during regeneration ^35^. Additionally, TGFB1 has been shown to be a key factor provided by stromal cells after injury to induce expression of fetal-reversion genes in facultative stem cells to aid in crypt recovery ^10^. Our data raises the question as to whether Paneth cells, in addition to stromal populations, secrete TGFB1 to support upper crypt cells after CBC loss. Nonetheless, the changes observed within the Paneth cells will require further experimentation to determine if these regenerative genes play a role in the crypt remodeling to adjust to CBC loss.

In addition to the profound importance for CBC maintenance at homeostasis, our study established the requirement for Paneth cell-derived DLL1 and DLL4 for regeneration after irradiation injury. Paneth cell or secretory dysfunction has long been associated with the development of inflammatory bowel disease and shown to exacerbate colitis-induced injury with delayed repair in mice ^23, 45–48^. However, functional studies involved DSS-induced colonic epithelial damage, and impaired regeneration was attributed primarily to microbial and immune cell dysregulation ^49^. Here, we showed that reduced DKO stem cell recovery after injury was independent of gut microbiota, with no evidence of inflammation, suggesting direct Paneth cell-to-stem cell signaling is involved in the regenerative response. This aligns with a previous study showing that intestinal epithelium-wide deletion of the Paneth cell-derived niche factor, Wnt3, impaired CBC expansion and crypt regeneration after injury from rotavirus infection, while proliferation at homeostasis was unaltered ^50^. Furthermore, we showed that DKO mice displayed significantly reduced organoid-forming efficiency, further implying that the impaired regeneration results from Paneth-stem cell interactions and is not mediated by immune or microbial populations. We also showed that OLFM4-expressing cells were depleted from DKO mice during crypt regeneration after injury, potentially implying that facultative stem cells driving regeneration failed to recover Notch signaling. These findings suggest that Paneth cells may provide Notch signal support to facultative and other stem cell populations in addition to CBCs during crypt regeneration.

Notably, crypt budding was not observed in DKO organoids, which may be due to lost Paneth cell - CBC interactions that are vital for crypt budding and crypt fission events ^51, 52^. Since we observed that expression of other niche components was normal in DKO Paneth cells, and the addition of recombinant DLL1 to the Matrigel rescued the impaired organoid-forming capability of DKO crypt cells, our data indicate that direct Paneth cell to CBC Notch signaling is crucial for stem cell recovery after injury.

In summary, our study showed that Paneth cell-specific ablation of DLL1 and DLL4 induced crypt cell remodeling, with depletion of CBCs and expansion of Paneth and endocrine cells within the crypt base. Crypts remodeled to support epithelial renewal; however, there was diminished overall stem cell function as demonstrated by reduced organoid forming efficiency and crypt regeneration after radiation injury. Altogether, we demonstrated the importance of the Paneth cell-specific Notch signal to maintain normal CBCs and appropriate levels of intestinal stem cell capacity to withstand injurious insults.

## Experimental Procedures

### Mice

Paneth cell-specific double Notch ligand knockout mice (DKO) were generated by crossing the following mouse strains: *Defa4^IRES-Cre^* ^53^, *ROSA26-LSL-tdTomato* (The Jackson Laboratory no. 007908), *Dll1^fl/fl^* ^54^, and *Dll4^fl/fl^* ^55^. An Inducible Paneth cell-specific ligand double knockout model (IDKO) was generated by crossing floxed-ligand mice with *Lyz1^3’UTR-IRES-CreERT2^*mice ^56^. Mice were housed in ventilated and automated watering cages with a 12 hour light cycle under specific pathogen-free conditions. Adult mice of both sexes, aged 2-5 months, were used for analyses. Protocols for the use of mice were approved by the University of Michigan Committee on the Use and Care of Animals.

### Animal Treatment Procedures

For analysis of proliferation, mice were given an intraperitoneal (i.p.) injection of 5-ethynyl-2’- deoxyuridine (EdU) (25 mg/kg) (Life Technologies) 1.5 h before tissue collection. For inducible Notch ligand ablation, IDKO mice and controls were injected with TAM daily (100 mg/kg i.p.) for 5 days with 2 days chase, or with a single TAM dose (200 mg/kg) followed by a 3-day chase period, as described in figure legends. For irradiation injury, mice were exposed to 12 Gy whole-body cesium irradiation with intestines harvested 3 days later. For microbiota depletion, mice were given broad-spectrum antibiotics in their drinking water as described ^41^: 100 U/L penicillin (Sigma-Aldrich, cat no P7793), 1g/L ampicillin (Sigma-Aldrich, cat no A9393), 1g/L neomycin (Sigma-Aldrich, cat no N1876), and 500 mg/L gentamycin (Sigma-Aldrich, cat no G1914). Mice were also gavaged every other day with 200 µL of 0.5 mg/mL metronidazole (Sigma-Aldrich, cat no M1547) and 1 mg/mL vancomycin (Cayman Chemical, cat no 15327) in sterile water. Antibiotic treatment was performed for two weeks prior to irradiation injury and continued until tissue harvest. Control and DKO antibiotic-treated mice were co-housed littermates, with bedding mixed among all cages 1 week prior to antibiotic treatment and irradiation injury.

### Histological Analysis

Intestinal tissue was fixed in 10% neutral buffered formalin overnight for paraffin embedding and tissue sections (4 µm) were stained as described ^15^. EdU-Click-iT kit (Life Technologies, cat. no. C10337) was used to identify proliferating cells. Hematoxylin & Eosin Stain kit (Vector Laboratories, Cat. no H-3502) was used for H&E staining. For immunofluorescence of paraffin-embedded tissues, antigen retrieval was performed with Antigen Unmasking Solution, Citrate-Based (Vector Laboratories, cat. no. H-3300-250). A buffer consisting of 0.01% Triton X, 5% BSA, and 10% goat serum was used for blocking (30 minutes-1hour) and for antibody dilution. Primary antibodies were incubated overnight at 4°C and secondary antibodies (1:400) were incubated at room temperature for 30 minutes-1 hour. The primary antibodies included the following: rabbit anti-OLFM4 (1:200, Cell Signal, cat. no. 39141S), rabbit anti-SLC12A2 (1:400, Abcam, cat. no. ab303518), rabbit anti-LYZ (1:750, DAKO, cat. no. A0099), rat anti-MMP7 (1:400, Vanderbilt, cat. no. 4334), rabbit anti-cleaved caspase-3 (1:50, Cell Signal, cat. no. 9664S), rabbit anti-MUC2 (1:200, Santa Cruz, cat. no. sc-15334), rabbit anti-CHGA (1:200, Abcam, cat. no. ab15160), rabbit anti-RFP (1:500, Rockland, cat no 600-401-379), mouse anti-RFP (1:400, Invitrogen, cat. no. MA5-15257), rabbit anti-DCAMKL1 (1:200, Abcam, cat. no. ab31704), mouse anti-E-cadherin (1:50, BD Biosciences, cat. no. 610181), mouse anti-CD45 (1:200, BD Pharmingen, Cat no 553076). Images were captured on a Nikon E800 microscope with Olympus DP controller software. Confocal images were captured on a Leica Stellaris SP5 inverted confocal microscope. Morphometrics were performed by counting positive or co-positive cells per well-oriented crypt, villus, or length of jejunum, as indicated in figure legends. Crypt depth measurements were performed using ImageJ software version 1.54p by measuring the length of crypts from H&E stained images. Brightness and contrast were altered for individual channels on most immunofluorescent images to maximize visibility, with alterations performed on the entire image with equal changes to control and DKO images. Red channels were changed from red to magenta in Adobe Photoshop 2026 to increase accessibility.

### In Situ Hybridization

Fluorescence in situ hybridization was performed on paraffin sections from intestinal tissue that was fixed for 24 hours in 10% neutral buffered formalin using the ACDbio RNAscope Multiplex Manual Assay v2 Kit (Advanced Cell Diagnostics, REF 323110) according to manufacturer’s instructions using Histo-clear II (National Diagnostics, 5989-27-5) for deparaffinization. For protease treatment, protease plus was applied for 30 minutes at 40 °C. The following probes were used: Mm-Ascl2-C1 (Cat No. 412211), Mm-Hmgcs2-C2 (Cat No. 437141), Mm-Olfm4-C2 (cat no 311831-C2), and mM-Tgfb1-C1 (cat No 407751). All incubations were performed at 40°C in a HybEZ hybridization system oven (ACD; 310010). After probe hybridization, slides were kept in 5X saline sodium citrate buffer overnight at 4 °C before amplification. For signal detection, TSA Cy5 (1:3500, Akoya Biosciences, a Quanterix company, Cat no NEL744001KT) or TSA Cy3 (1:3000, Akoya Biosciences, a Quanterix company, Cat no NEL745E001KT) were diluted in the kit-provided TSA buffer. Primary antibody incubation of mouse anti-CDH1 (1:50, BD Biosciences, cat. no. 610181) and/or rabbit anti-Lysozyme (1:500, DAKO, cat. no. A0099) diluted in 0.03% Triton X, 20% goat serum, 1X PBS was performed after signal amplification of RNAscope probes.

### Small Intestinal Crypt and Single Cell Isolation

To isolate crypts, the first 10 cm of jejunum was removed, cut into 2 cm pieces, opened longitudinally, and washed with phosphate-buffered saline (DPBS; Gibco, REF 14190-144, LOT 3062464) on ice to remove luminal contents, rinsed in DPBS supplemented with 1X Pen/Strep (Gibco, Cat no 15140122), 1X Gentamycin (Gibco, Cat no 15750-060), and 1 μg/mL Fungizone (R&D Systems, Cat no B23192), and scraped gently with a sterile glass slide to remove villi and debris. To isolate crypts, scraped tissue was incubated in 10 mL of 15 mM ethylenediaminetetraacetic acid (EDTA, pH 8.0, Invitrogen REF AM9260G, LOT 3171453) in DPBS with antibiotics, rocking gently on ice for 35 minutes, and washed with cold DPBS, followed by vortexing at maximum speed for 2 minutes. The resulting crypt solution was filtered through a 70 µm strainer to isolate crypts.

For single-cell preparation, DNase I (100 µg/mL Sigma Aldrich, REF 101041590001, LOT 85201300) in 20% FBS (Corning, Cat no 35-070-CV) in 5 mL 1X HBSS (Gibco, Hanks’ Balanced Salt Solution, REF 14175-095) was added to isolated crypts. Crypts were pelleted and then dissociated in Dispase II (Sigma Aldrich, Cat no 04942078001) at 1U/mL diluted in 5mL HBSS, incubated for 10 minutes at 37°C with intermittent shaking by hand. Single cells were pelleted at 1280 x g for 5 minutes at 4°C and resuspended in 1X HBSS containing 100 µg/mL DNase I and 20% FBS, filtered through a 40 µm cell strainer, pelleted, and resuspended in either 1mL of DMEM F12 (Gibco REF 12634-010, LOT 3086148) with antibiotics for organoid culture or 0.5 mL flow media (20% FBS in DMEM/F12 with 10 µM Y-27632 dihydrochloride (Tocris Bioscience, Cat no 1254) for Paneth cell isolation.

### Organoid Culture

Isolated crypts were diluted with DMEM/F12 containing antibiotics, centrifuged at 500 x g for 6 minutes at 4°C, and resuspended in Corning Matrigel Matrix (Corning, cat no 356255) and plated as 100-200 crypts per 10 µL patty in 48 well plates. Crypts were counted via hemocytometer. After incubation at 37°C for 15–30 minutes, Matrigel patties were overlaid with 200 µL of culture media. Organoid high-WNT growth media contained 50% L-WRN conditioned media (University of Michigan Human Organoid Core), 20% FBS, 1X HEPES (Gibco, Cat no 15630130), 1X GlutaMAX (Gibco, Cat no 35050-061) with antibiotics and Y-27632 as listed above in DMEM/F12 (Gibco Cat 11320033). Differentiation media contained 10% R-Spondin-1 conditioned media (University of Michigan TTML Core), Noggin (100 ng/ml, R&D Systems, Cat no 6057-NG), EGF (50 ng/ml, R&D Systems, Cat no 236-EG), HEPES, GlutaMAX, B27 (Gibco, Cat no 17504001), with antibiotics and Y-27632 in DMEM/F12.

### Organoid Forming Efficiency

Crypts were dissociated into single cells and plated at a dilution of 12,000 cells per 10 µL of Matrigel seeded in 48-well plates and organoid numbers were counted 5 days after culture in organoid growth medium. Cells were counted via hemocytometer. Organoid-forming efficiency was calculated using the following formula: number of organoids formed/ number of cells plated x 100. Organoids were counted in a blinded manner with 6-8 wells counted per mouse. For Notch ligand supplementation, DLL1 (1 µM, ACRO Biosystems, Cat. # DL1-H52H8) was added directly to the Matrigel immediately prior to seeding 10,000 cells per 10 µL of Matrigel, as previously described ^12^. Well-view pictures were taken on a dissecting/stereo scope (Olympus SZX16) and organoid-view pictures were taken on a Nikon E800 microscope or an Olympus IX71 inverted-fluorescent microscope.

### FACS Isolation of Paneth cells and RNA-sequencing

Two male and two female mice from each genotype were used for Paneth cell transcriptomic analysis. To isolate Paneth cells, Y-28623 (10 µM) was added at the beginning of crypt isolation and maintained throughout each step to increase cell viability. After single-cell dissociation, cells were stained with EpCAM- SuperBright645 (1:200, Invitrogen, Cat. no 64-5791-82) for 30 minutes on ice and washed twice with 20% FBS in DMEM/F12 and isolated with a Thermo Fisher Scientific Bigfoot Spectral Cell Sorter (University of Michigan Flow cytometry core). DRAQ7 (1:500, Invitrogen, Cat. no D15106) was added 10 minutes prior to sorting to detect cell viability. Epcam and tdTomato double-positive Paneth cells (40,000) were sorted into 300 µL of RLT lysis buffer (Qiagen, cat no 74004) containing β-mercaptoethanol (Sigma Aldrich, cat no M3148) per manufacturer instructions, and vortexed at maximum speed before flash freezing in liquid N_2_.

Paneth cell RNA was isolated using the RNeasy micro kit (Qiagen, cat no 74004) according to manufacturers’ instructions. RNA quality and concentration were confirmed by the University of Michigan Advanced Genomics Core using the Agilent Bioanalyzer and Invitrogen Qubit Fluorometer. An Illumina low-input poly (A) enrichment library prep and next-generation sequencing were carried out in the Advanced Genomics Core at the University of Michigan on the Illumina NovaSeq X with 10B lane sequencing, 300 cycle to a depth of ∼60 million reads. Sequencing data were transferred to the University of Michigan Great Lakes high-performance computing cluster for analysis. The nf-core/rnaseq (v3.19.0-g0bb032c) Nextflow pipline was used for data processing and alignment. Sequencing read quality was assessed via FastQC and MultiQC. Reads were aligned to the Mus musculus genome build GRCm38 using STAR v2.7.10a. Aligned reads were assigned to genes using RSEM v1.3.1. Differential gene expression was calculated as previously described ^57^. Briefly, read count matrices were imported into R/Bioconductor (version 4.5.1) package edgeR (version 4.6.3). Genes with low expression were excluded from further analysis using the edgeR *filterByExpr*() function with default settings. Normalization factors and dispersion were calculated prior to generating a log2-transformed normalized counts per million (CPM) matrix for downstream analysis. Differential gene expression between each treatment (average of four replicates) and the relevant controls (average of four replicates) was calculated using quasi-likelihood negative binomial generalized log-linear modeling in edgeR. Genes were considered differentially expressed between treatment and control when both the false discovery rate (FDR) and the adjusted p-value were below 0.05. Genes with negative average log2CPM were removed from analysis. Multidimensional scaling dimension reduction analysis and volcano plot generation were performed in R/Bioconductor (version 4.5.1). Heatmaps were generated in GraphPad prism from Z scores based on normalized log2CPM values calculated, as described above, in R/Bioconductor. The list of 100 upregulated genes, ranked by F score, was uploaded to Metascape.org to generate top GO terms. The −log10(Padj) scores and GO terms generated by Metascape were used to create a horizontal bar graph in GraphPad Prism. Flow plots were generated using FlowJo software version 10.10.0.

### Measurement of mRNA abundance by RT-qPCR

Isolated crypts were flash frozen in RLT lysis buffer (Qiagen, cat. no. 74104), thawed on ice, homogenized for 15 seconds with a Polytron homogenizer, and RNA was isolated using the RNeasy Mini kit (Qiagen, cat. no. 74104) with DNase I treatment as per manufacturer instructions. RNA quality and concentration were measured by Bioanalyzer and Qubit by the University of Michigan Advanced Genomics Core. Only crypt RNA samples with RIN > 8 were used. mRNA abundance was measured by RT-qPCR relative to *Gapdh* and expressed as fold change (2 ^−ΔΔCt^).

### Statistical Analysis

Prism software (GraphPad) was used for statistical analysis. All statistical comparisons were performed with three to eight biological replicates per group, as indicated in figure legends. Quantitative data are presented as mean ± SD. Comparisons were analyzed between two groups with an unpaired two-tailed Student’s t test with Welch’s correction used for t-tests whenever the variances of the groups significantly differed. For analysis of organoid-forming efficiency with controls and DKO in vehicle- vs DLL1- supplemented Matrigel, we used two-way ANOVA with Bonferroni’s correction since two independent variables were tested. Significance is reported as ^∗^p < 0.05, ^∗∗^p < 0.01, ^∗∗∗^p < 0.001, ^∗∗∗∗^p < 0.0001. For transcriptomic analyses, four mice per group were used. Both an FDR of less than 0.05 and an adjusted p-value of less than 0.05 were used to determine the significance of differentially expressed genes.

## Resource Availability

The bulk RNA-seq data have been deposited at Gene Expression Omnibus (GEO) at GEO: GSE344487. Any additional information required to reanalyze the data reported in this paper is available from the corresponding author upon request.

## Supporting information

Document S1 including all supplemental figures (S1-S6)

## Acknowledgements

We thank the Jason Spence and Kate Walton labs for use of their RNAscope mRNA probes and the Yatrik Shah lab for use of antibiotic reagents and their guidance on antibiotic treatment of mice. We also thank the Nataliya Razumilava lab for use of reagents and Katie Patton for mouse colony maintenance.

## Grant Support

This project was funded by NIH R01 DK118023 (LCS), NIH R01 ES022802 (JAC), and NIH F32 1F32DK142253 (MQ).

## Author Contributions

L.C.S. and M.Q. conceptualized the project with subsequent contributions from P.J.D. The majority of the investigation and analysis was performed by M.Q. with contributions from T.M.K and J.C. Key resources were provided by P.J.D. and N.G. The manuscript was written by M.Q. and L.C.S. All authors reviewed the manuscript and provided critical feedback. L.C.S. provided supervision and obtained funding.

## Declaration of Interests

The authors declare no competing interests.

## Abbreviations

AB/PAS: Alcian blue/ periodic acid-Schiff
CBC: crypt base columnar cell
CC3: cleaved caspase 3
CDH1: epithelial cadherin
CHGA: chromogranin A
DCLK1: doublecortin-like kinase 1
DEGs: differentially expressed genes
DKO: Notch ligand-depleted mice (*Dll1^fl/fl^*’; *Dll4^fl/fl^*; *Defa4^IRES-Cre^*; *ROSA26-LSL- tdTomato*)
DLL1: delta-like ligand 1
DLL4: delta-like ligand 4
EdU: 5-ethynyl-2’-deoxyuridine
IDKO: inducible Notch ligand-depleted mice (*Dll1^fl/fl^*; *Dll4^fl/fl^*; *Lyz1^3’UTR-IRES-CreERT2^*)
GO: gene ontology
LYZ: lysozyme
MDS: multidimensional scaling
MMP7: matrix metalloproteinase-7
MUC2: mucin-2
OLFM4: olfactomedin 4
RNA-seq: RNA-sequencing
RT-qPCR: reverse transcriptase quantitative polymerase chain reaction
SLC12A2: solute carrier family 12 member 2
TAM: tamoxifen.

---

**Table 1.**
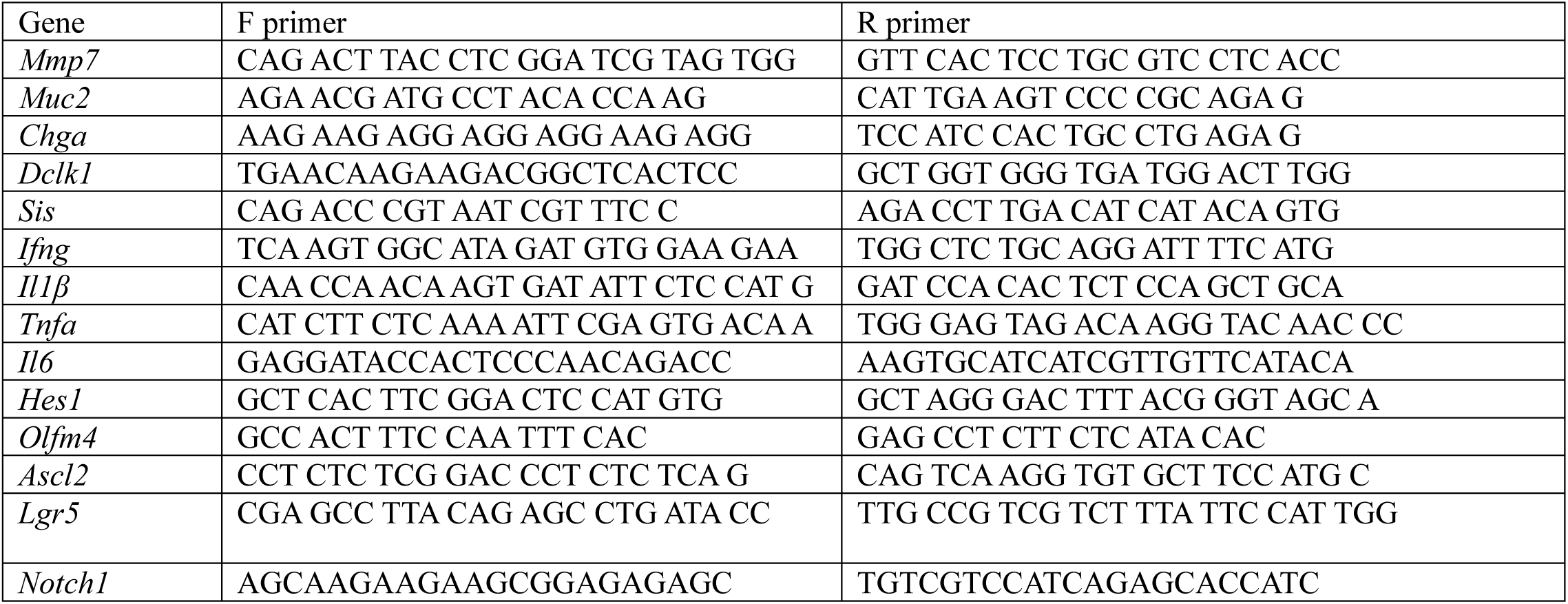
RT-qPCR primers.

## Supplemental Information

Document S1. Figures S1-S6

Table S1. Key unchanged Paneth cell genes, Excel file, related to Figure 4

Genes listed by their description, FDR > 0.05 is considered unchanged.

Table S2. DEGs in DKO Paneth cells, Excel file, related to Figure 4

DEGs ordered by significance. FDR < 0.05 is considered differentially expressed. Log2FC > 0 is considered upregulated. Log2FC < 0 is considered downregulated.

Table S3. GO term analysis of top 100 upregulated DEGs.

Full list of GO terms in order of significance by log10(P) from top 100 upregulated genes ranked by F

Table S4. GO term analysis of downregulated DEGs.

Full list of GO terms in order of significance by log10(P) from all downregulated genes ranked by F

## Notes

https://www.ncbi.nlm.nih.gov/geo/query/acc.cgi?acc=GSE344487

### Competing Interest Statement

The authors have declared no competing interest.

## References

1. Barker N, van Es JH, Kuipers J, et al. Identification of stem cells in small intestine and colon by marker gene Lgr5. Nature 2007;449:1003–7.

2. Tetteh PW, Basak O, Farin HF, et al. Replacement of Lost Lgr5-Positive Stem Cells through Plasticity of Their Enterocyte-Lineage Daughters. Cell Stem Cell 2016;18:203–13.

3. Jones JC, Brindley CD, Elder NH, et al. Cellular Plasticity of Defa4(Cre)-Expressing Paneth Cells in Response to Notch Activation and Intestinal Injury. Cell Mol Gastroenterol Hepatol 2019;7:533–554.

4. Schmitt M, Schewe M, Sacchetti A, et al. Paneth Cells Respond to Inflammation and Contribute to Tissue Regeneration by Acquiring Stem-like Features through SCF/c-Kit Signaling. Cell Rep 2018;24:2312–2328 e7.

5. Yu S, Tong K, Zhao Y, et al. Paneth Cell Multipotency Induced by Notch Activation following Injury. Cell Stem Cell 2018;23:46–59 e5.

6. van Es JH, Sato T, van de Wetering M, et al. Dll1+ secretory progenitor cells revert to stem cells upon crypt damage. Nat Cell Biol 2012;14:1099–1104.

7. Yan KS, Gevaert O, Zheng GXY, et al. Intestinal Enteroendocrine Lineage Cells Possess Homeostatic and Injury-Inducible Stem Cell Activity. Cell Stem Cell 2017;21:78–90 e6.

8. Yui S, Azzolin L, Maimets M, et al. YAP/TAZ-Dependent Reprogramming of Colonic Epithelium Links ECM Remodeling to Tissue Regeneration. Cell Stem Cell 2018;22:35–49 e7.

9. Nusse YM, Savage AK, Marangoni P, et al. Parasitic helminths induce fetal-like reversion in the intestinal stem cell niche. Nature 2018;559:109–113.

10. Chen L, Qiu X, Dupre A, et al. TGFB1 induces fetal reprogramming and enhances intestinal regeneration. Cell Stem Cell 2023;30:1520–1537 e8.

11. Meyer AR, Brown ME, McGrath PS, et al. Injury-Induced Cellular Plasticity Drives Intestinal Regeneration. Cell Mol Gastroenterol Hepatol 2022;13:843–856.

12. VanDussen KL, Carulli AJ, Keeley TM, et al. Notch signaling modulates proliferation and differentiation of intestinal crypt base columnar stem cells. Development 2012;139:488–97.

13. van Es JH, van Gijn ME, Riccio O, et al. Notch/gamma-secretase inhibition turns proliferative cells in intestinal crypts and adenomas into goblet cells. Nature 2005;435:959–63.

14. Milano J, McKay J, Dagenais C, et al. Modulation of notch processing by gamma-secretase inhibitors causes intestinal goblet cell metaplasia and induction of genes known to specify gut secretory lineage differentiation. Toxicol Sci 2004;82:341–58.

15. Bohin N, Keeley TM, Carulli AJ, et al. Rapid Crypt Cell Remodeling Regenerates the Intestinal Stem Cell Niche after Notch Inhibition. Stem Cell Reports 2020;15:156–170.

16. Tsai YH, VanDussen KL, Sawey ET, et al. ADAM10 regulates Notch function in intestinal stem cells of mice. Gastroenterology 2014;147:822–834 e13.

17. Carulli AJ, Keeley TM, Demitrack ES, et al. Notch receptor regulation of intestinal stem cell homeostasis and crypt regeneration. Dev Biol 2015;402:98–108.

18. Shaya O, Binshtok U, Hersch M, et al. Cell-Cell Contact Area Affects Notch Signaling and Notch-Dependent Patterning. Dev Cell 2017;40:505–511 e6.

19. Farin HF, Van Es JH, Clevers H. Redundant sources of Wnt regulate intestinal stem cells and promote formation of Paneth cells. Gastroenterology 2012;143:1518–1529 e7.

20. Sato T, van Es JH, Snippert HJ, et al. Paneth cells constitute the niche for Lgr5 stem cells in intestinal crypts. Nature 2011;469:415–8.

21. Pellegrinet L, Rodilla V, Liu Z, et al. Dll1- and dll4-mediated notch signaling are required for homeostasis of intestinal stem cells. Gastroenterology 2011;140:1230–1240 e1-7.

22. Durand A, Donahue B, Peignon G, et al. Functional intestinal stem cells after Paneth cell ablation induced by the loss of transcription factor Math1 (Atoh1). Proc Natl Acad Sci U S A 2012;109:8965–70.

23. Quintero M, Liu S, Xia Y, et al. Cdk5rap3 is essential for intestinal Paneth cell development and maintenance. Cell Death Dis 2021;12:131.

24. Kim TH, Escudero S, Shivdasani RA. Intact function of Lgr5 receptor-expressing intestinal stem cells in the absence of Paneth cells. Proc Natl Acad Sci U S A 2012;109:3932–7.

25. van Es JH, Wiebrands K, Lopez-Iglesias C, et al. Enteroendocrine and tuft cells support Lgr5 stem cells on Paneth cell depletion. Proc Natl Acad Sci U S A 2019;116:26599–26605.

26. Cheng CW, Biton M, Haber AL, et al. Ketone Body Signaling Mediates Intestinal Stem Cell Homeostasis and Adaptation to Diet. Cell 2019;178:1115–1131 e15.

27. Dekaney CM, King S, Sheahan B, et al. Mist1 Expression Is Required for Paneth Cell Maturation. Cell Mol Gastroenterol Hepatol 2019;8:549–560.

28. Bhattacharya S, Tie G, Singh PNP, et al. Intestinal secretory differentiation reflects niche-driven phenotypic and epigenetic plasticity of a common signal-responsive terminal cell. Cell Stem Cell 2025;32:952–969 e8.

29. Shroyer NF, Wallis D, Venken KJ, et al. Gfi1 functions downstream of Math1 to control intestinal secretory cell subtype allocation and differentiation. Genes Dev 2005;19:2412–7.

30. Viragova S, Li D, Klein OD. Activation of fetal-like molecular programs during regeneration in the intestine and beyond. Cell Stem Cell 2024;31:949–960.

31. Zhang J, Ye, J., Ren, Y., Zuo, J. Dai, W., He, Y., Tan, M., Song, W., Yuan Y. Intracellular activation of complement C3 in Paneth cells improves repair of intestinal epithelia during acute injury. Immunotherapy 2018;10:1325–1336.

32. Ye J, Yuan K, Dai W, et al. The mTORC1 signaling modulated by intracellular C3 activation in Paneth cells promotes intestinal epithelial regeneration during acute injury. Int Immunopharmacol 2019;67:54–61.

33. Schraufstatter IU, Khaldoyanidi SK, DiScipio RG. Complement activation in the context of stem cells and tissue repair. World J Stem Cells 2015;7:1090–108.

34. Hu DJ, Yun J, Elstrott J, et al. Non-canonical Wnt signaling promotes directed migration of intestinal stem cells to sites of injury. Nat Commun 2021;12:7150.

35. Cheng J, Wu H, Cui Y. WNT4 promotes the symmetric fission of crypt in radiation-induced intestinal epithelial regeneration. Cell Mol Biol Lett 2024;29:158.

36. Takashima S, Sharma R, Chang W, et al. STAT1 regulates immune-mediated intestinal stem cell proliferation and epithelial regeneration. Nat Commun 2025;16:138.

37. Shindo R, Katagiri T, Komazawa-Sakon S, et al. Regenerating islet-derived protein (Reg)3beta plays a crucial role in attenuation of ileitis and colitis in mice. Biochem Biophys Rep 2020;21:100738.

38. McMahan RH, Najarro KM, Giesy LE, et al. Intestinal REG3G Protects Against Gastrointestinal Dysfunction in a Murine Model of Ethanol Intoxication and Burn Injury. Shock 2026;65:528–537.

39. Bhattacharyya S, Fang F, Tourtellotte W, et al. Egr-1: new conductor for the tissue repair orchestra directs harmony (regeneration) or cacophony (fibrosis). J Pathol 2013;229:286–97.

40. Bohin N, McGowan KP, Keeley TM, et al. Insulin-like Growth Factor-1 and mTORC1 Signaling Promote the Intestinal Regenerative Response After Irradiation Injury. Cell Mol Gastroenterol Hepatol 2020;10:797–810.

41. Das NK, Schwartz AJ, Barthel G, et al. Microbial Metabolite Signaling Is Required for Systemic Iron Homeostasis. Cell Metab 2020;31:115–130 e6.

42. Annunziata F, Rasa SMM, Krepelova A, et al. Paneth cells drive intestinal stem cell competition and clonality in aging and calorie restriction. Eur J Cell Biol 2022;101:151282.

43. Capdevila C, Miller J, Cheng L, et al. Time-resolved fate mapping identifies the intestinal upper crypt zone as an origin of Lgr5+ crypt base columnar cells. Cell 2024;187:3039–3055 e14.

44. Malagola E, Vasciaveo A, Ochiai Y, et al. Isthmus progenitor cells contribute to homeostatic cellular turnover and support regeneration following intestinal injury. Cell 2024;187:3056–3071 e17.

45. Stappenbeck TS, McGovern DPB. Paneth Cell Alterations in the Development and Phenotype of Crohn’s Disease. Gastroenterology 2017;152:322–326.

46. Cadwell K, Liu JY, Brown SL, et al. A key role for autophagy and the autophagy gene Atg16l1 in mouse and human intestinal Paneth cells. Nature 2008;456:259–63.

47. Kaser A, Lee AH, Franke A, et al. XBP1 links ER stress to intestinal inflammation and confers genetic risk for human inflammatory bowel disease. Cell 2008;134:743–56.

48. Cai Y, Zhu G, Liu S, et al. Indispensable role of the Ubiquitin-fold modifier 1-specific E3 ligase in maintaining intestinal homeostasis and controlling gut inflammation. Cell Discov 2019;5:7.

49. Liu TC, Gurram B, Baldridge MT, et al. Paneth cell defects in Crohn’s disease patients promote dysbiosis. JCI Insight 2016;1:e86907.

50. Zou WY, Blutt SE, Zeng XL, et al. Epithelial WNT Ligands Are Essential Drivers of Intestinal Stem Cell Activation. Cell Rep 2018;22:1003–1015.

51. Pin C, Parker A, Gunning AP, et al. An individual based computational model of intestinal crypt fission and its application to predicting unrestrictive growth of the intestinal epithelium. Integr Biol (Camb) 2015;7:213–28.

52. Langlands AJ, Almet AA, Appleton PL, et al. Paneth Cell-Rich Regions Separated by a Cluster of Lgr5+ Cells Initiate Crypt Fission in the Intestinal Stem Cell Niche. PLoS Biol 2016;14:e1002491.

53. Burger E, Araujo A, Lopez-Yglesias A, et al. Loss of Paneth Cell Autophagy Causes Acute Susceptibility to Toxoplasma gondii-Mediated Inflammation. Cell Host Microbe 2018;23:177–190 e4.

54. Hozumi K, Negishi N, Suzuki D, et al. Delta-like 1 is necessary for the generation of marginal zone B cells but not T cells in vivo. Nat Immunol 2004;5:638–44.

55. Koch U, Fiorini E, Benedito R, et al. Delta-like 4 is the essential, nonredundant ligand for Notch1 during thymic T cell lineage commitment. J Exp Med 2008;205:2515–23.

56. Balasubramanian I, Bandyopadhyay S, Flores J, et al. Infection and inflammation stimulate expansion of a CD74(+) Paneth cell subset to regulate disease progression. EMBO J 2023;42:e113975.

57. Sala-Hamrick KE, Tapaswi A, Polemi KM, et al. High-Throughput Transcriptomics of Nontumorigenic Breast Cells Exposed to Environmentally Relevant Chemicals. Environ Health Perspect 2024;132:47002.

