## Supplementary material for "Paneth cell-derived Notch ligands DLL1 and DLL4 are crucial for intestinal crypt base stem cells": Document S1 including all supplemental figures (S1-S6)

Samuelson

Figures S1-S6.

Figure S1

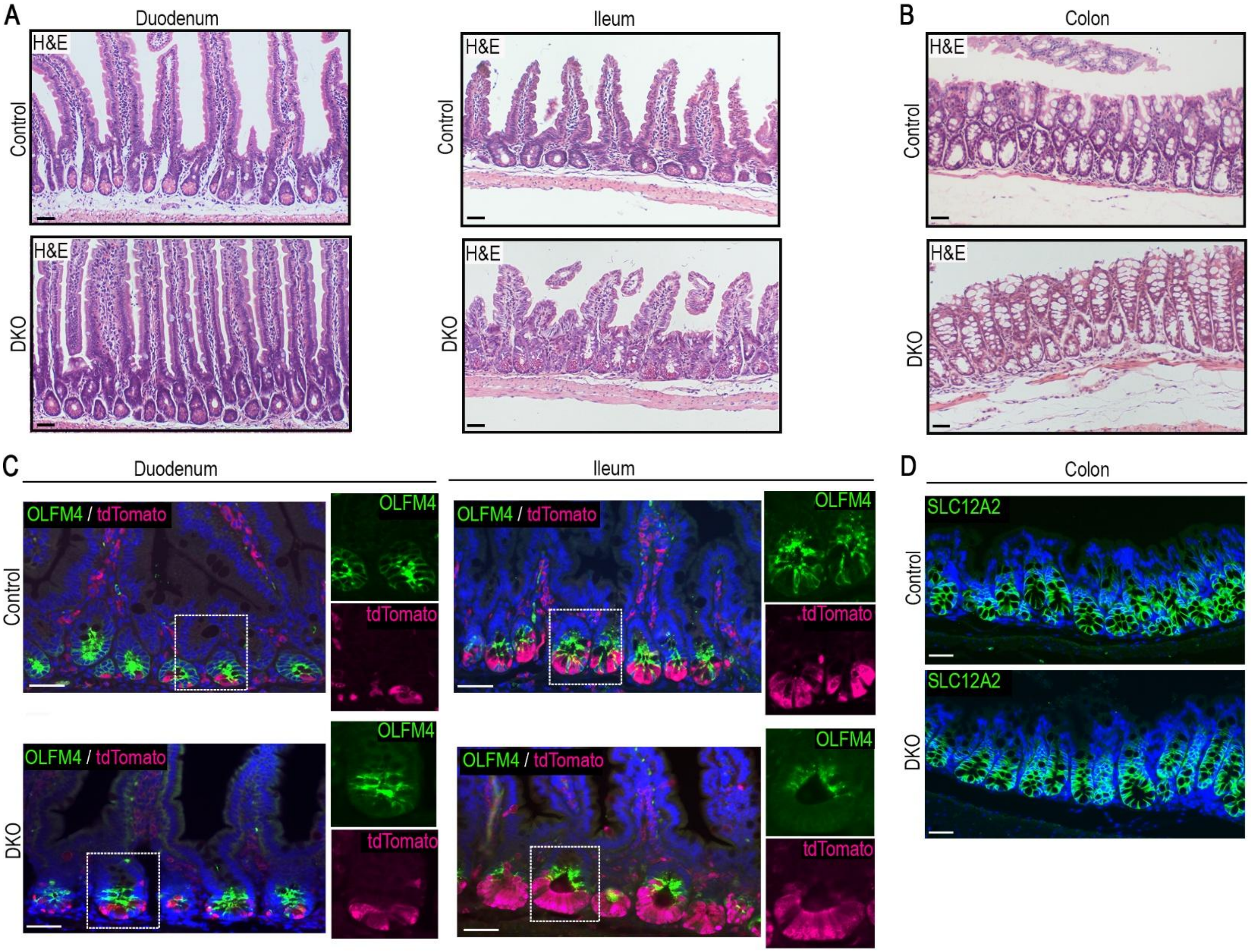


<u>Figure S1. Effect of Paneth cell-specific Notch ligand ablation on different regions of the intestine.</u>

(A) H&E staining of the duodenum and ileum from DKO (*$Dll1^{f/f}$; $Dll4^{f/f}$; $Defa4^{IRES\text{-}Cre}$; ROSA26-LSL-tdTomato*) and control mice. Scale bar, 50 µm.
(B) H&E staining of the DKO mouse colon. Scale bar, 50 µm.
(C) IF staining for the CBC marker OLFM4 (green) and the tdTomato reporter (magenta) with DAPI (blue) in the duodenum and ileum, marking CBCs and recombination activity of *$Defa4^{IRES\text{-}Cre}$*. Scale bar, 50 µm.
(D) IF staining for SLC12A2 (green) with DAPI (blue) showing normal CBC localization in the colon. Scale bar, 50 µm.

Figure S2

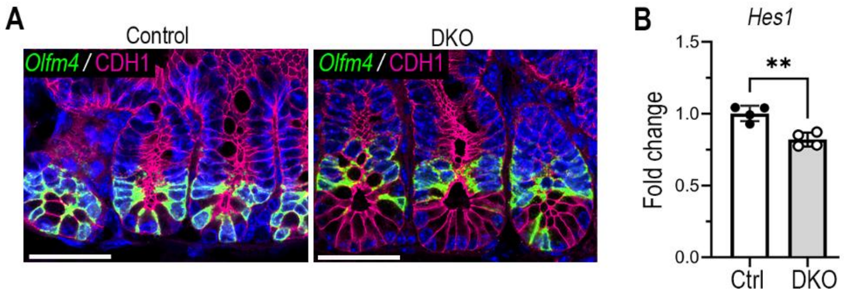


<u>Figure S2. Effect of Paneth cell-specific Notch ligand ablation on crypt cell Notch activity</u>
(A) In situ hybridization for the CBC marker and Notch target gene *Olfm4* (green) by RNAscope and IF staining for E-cadherin (CDH1, magenta) with DAPI (blue). Scale bar, 50 µm.
(B) RT-qPCR measurement of the abundance of the Notch target gene *Hes1* in RNA isolated from jejunal crypts. Quantitative data are presented as mean fold-change ± SD (Ctrl versus DKO by unpaired two-tailed Student's t-test; **p < 0.001, n = 4 mice/group).

Figure S3

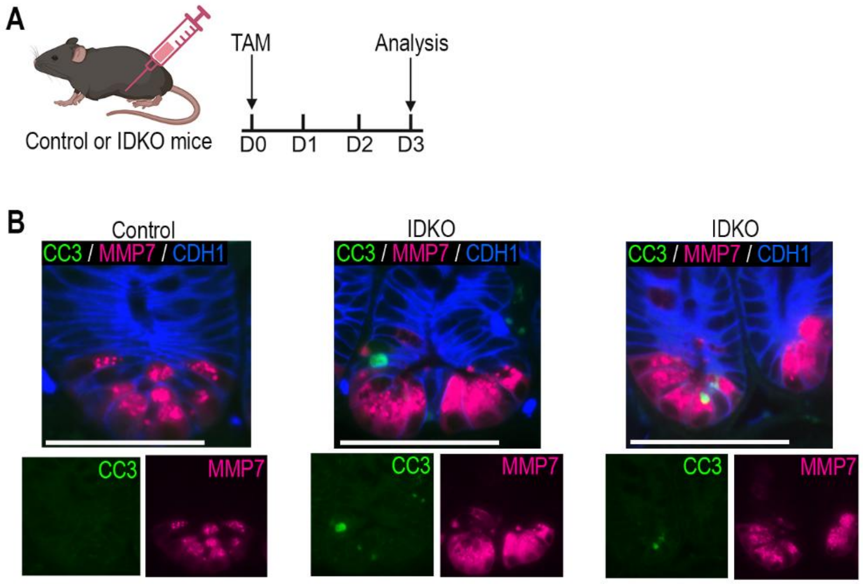


Figure S3. Apoptotic crypt cells are present 3 days after inducing Paneth cell-specific Notch ligand ablation.
(A) IDKO mice (*Lyz1*$^{3'UTR\text{-}IRES\text{-}CreERT2}$; *Dll1*$^{f/f}$; *Dll4*$^{f/f}$) and control mice were treated with a single dose of 200 mg/kg of Tamoxifen (TAM) with intestines harvested 3 days later.
(B) IF co-staining for the cell death marker cleaved caspase 3 (CC3, green), the Paneth cell marker MMP7 (magenta), and CDH1 (blue) in jejunal sections. Shown are high-powered views of crypts from control or IDKO mice, with outsets showing individual channels for CC3 (green) or MMP7 (magenta). Scale bars, 50 µm.

Figure S4

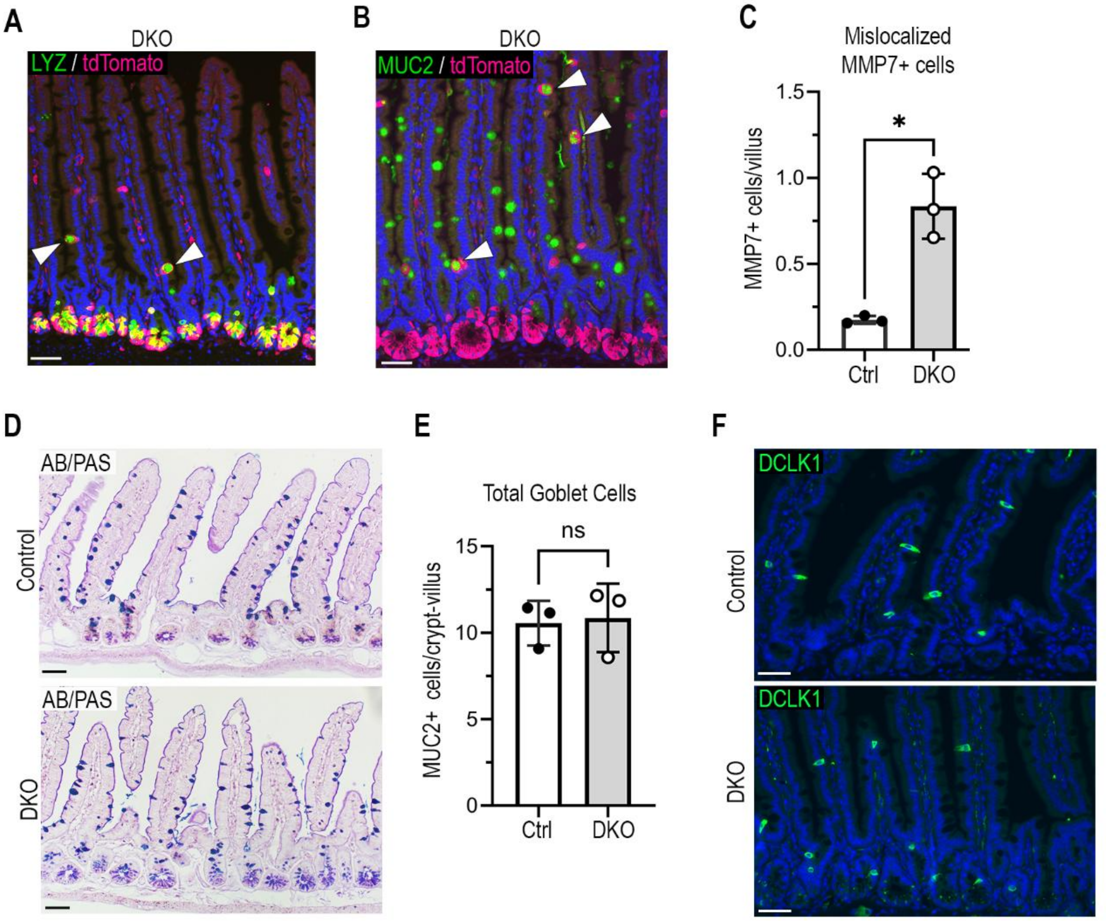


Figure S4. Effect of Paneth cell-specific Notch ligand ablation on intestinal epithelial cell differentiation

(A) Co-IF staining for the Paneth cell marker lysozyme (LYZ, green) and tdTomato (magenta) in DKO jejunal sections with DAPI (blue). Arrowheads point to co-labeled cells. Scale bar, 50 µm.

(B) Co-IF staining for the goblet cell marker MUC2 (green) and tdTomato (magenta) in DKO jejunal sections. Arrows point to co-labeled cells. Scale bar, 50 µm.

(C) Quantification of $MMP7^{+}$ Paneth-like cells mislocalized to the villus region of the DKO jejunum. All $MMP7^{+}$ cells in well-oriented villi within a 4cm strip of jejunum were counted per mouse (100-200 villi). The numbers of $MMP7^{+}$ cells are presented as mean ± SD (*$p < 0.01$, Ctrl versus DKO by unpaired two-tailed Welch's t-test; n = 3 mice/group).

(D) Goblet cells were stained with Alcian Blue and Periodic Acid Schiff (AB/PAS) in jejunal sections. Scale bar, 50 µm.

(E) Quantification of $MUC2^{+}$ goblet cells in DKO jejunum. All $MUC^{+}$ cells for all well-oriented villus-crypt units within a 4 cm strip of jejunum were counted per mouse (100-200 villi/crypts). The numbers of $MUC2^{+}$ cells are presented as mean ± SD (*$p < 0.01$, Crtl versus DKO by unpaired two-tailed Student's t-test; n = 3 mice/group).

(F) IF staining for the tuft cell marker DCLK1 (green) with DAPI (blue) shows apparently normal tuft cell differentiation in the DKO jejunum. Scale bar, 50 µm

Figure S5

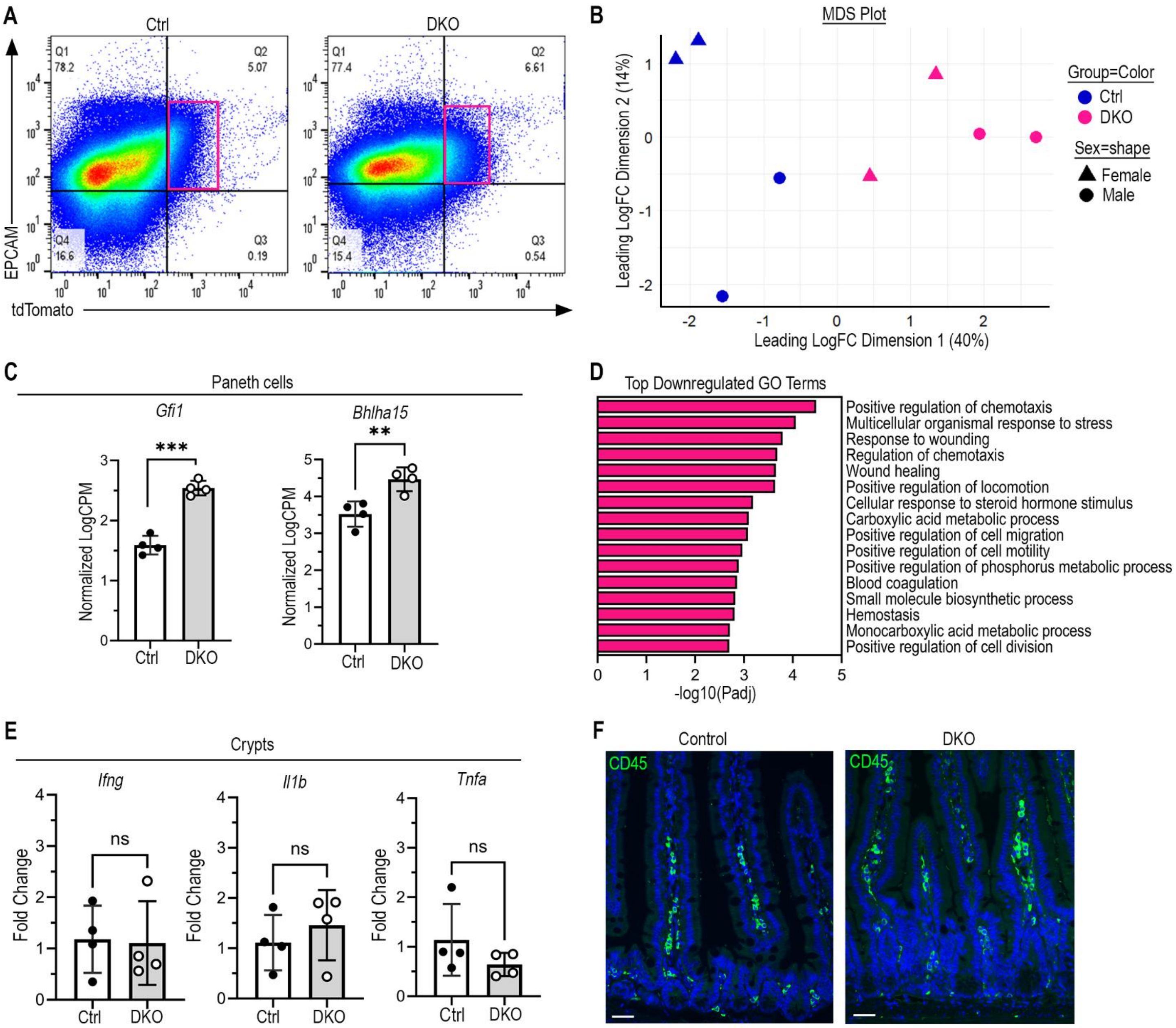


Figure S5. RNAseq analysis of sorted Paneth cells from control and DKO mice

(A) Representative flow plots from control and DKO mice showing cellular staining intensity for the epithelial cell marker EPCAM and the Paneth cell marker tdTomato generated with FlowJo software. Double-labeled EPCAM$^{+}$ / TdTomato$^{+}$ Paneth cells were isolated from Ctrl and DKO mice (n = 4 mice/group, 2 females and 2 males) for bulk RNAseq analysis.

(B) Multidimensional scaling (MDS) plot showing separation of DKO and control samples. Ctrl or DKO group is shown by color and sex is indicated by shape (n = 4 mice, group)

(C) Differential expression of genes involved in Paneth cell development in Ctrl and DKO mice. Data is shown as mean normalized counts per million (log2CPM) ± SD (Ctrl versus DKO by unpaired two-tailed Student's t-test; ***p < 0.0001, **p < 0.001, n = 4 mice/group).

(D) Top GO terms for all downregulated DEG between Ctrl and DKO. The list of downregulated genes, ranked by F score was uploaded to Metascape.org to generate top GO terms. The -log10(Padj) scores and GO terms generated by Metascape were used to create a horizontal bar graph in GraphPad Prism. Full list of GO terms shown in Table S4.

(E) Measurement of mRNA abundance of damage-associated inflammatory markers by RT-qPCR analysis of RNA isolated from jejunal crypts. Data are displayed as mean fold-change ± SD (Ctrl versus DKO by unpaired two-tailed Student's t-test or Welch's t-test when variance between DKO and controls differed; ns, not significant; n = 4 mice/group).

(F) IF staining for the pan-immune cell marker, CD45 (green) with DAPI (blue) in jejunal sections. Scale bars, 50 μm

Figure S6

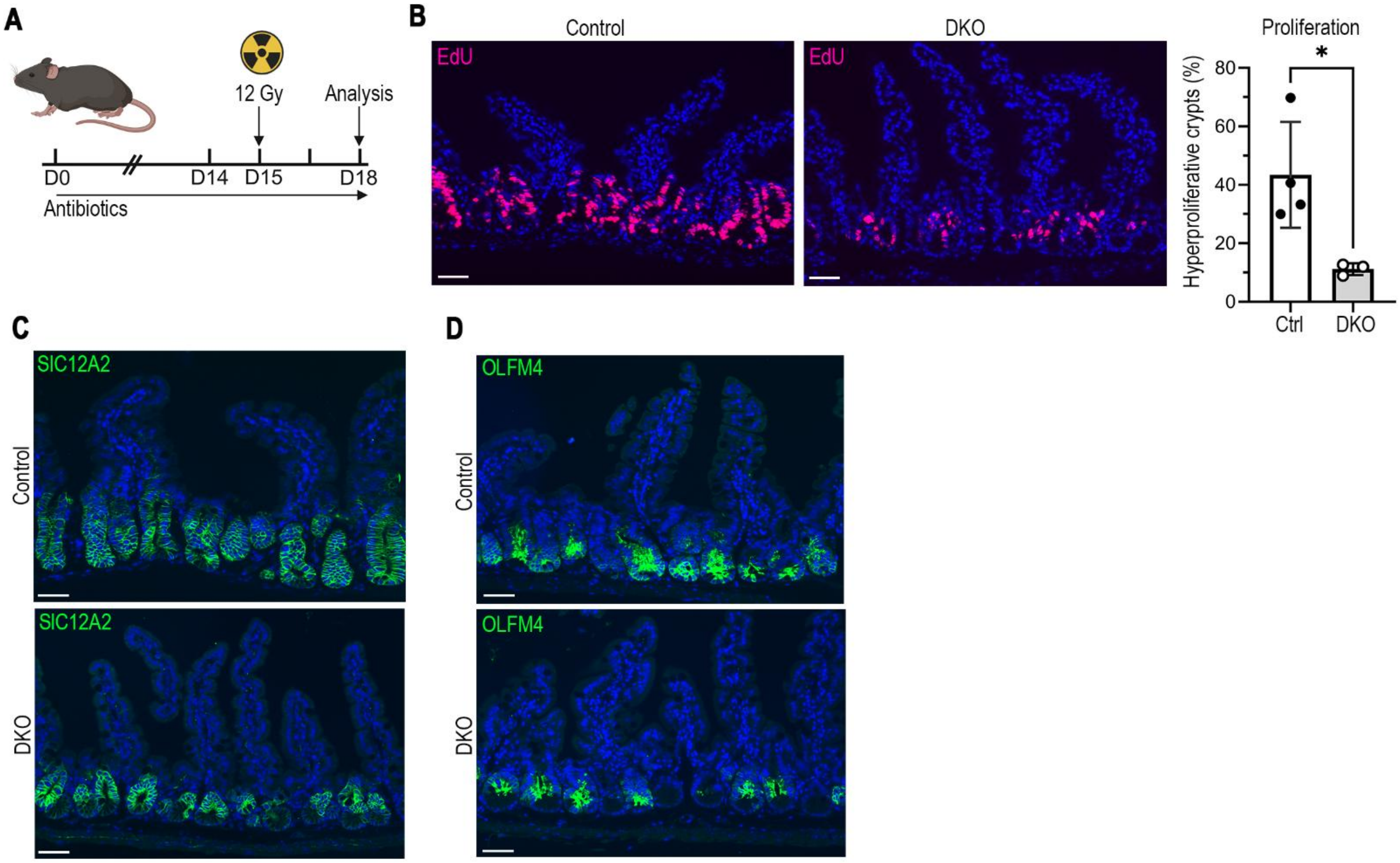


<u>Figure S6. Impaired crypt regeneration after radiation injury in DKO mice after microbial depletion</u>

(A) Experimental schematic. Mice were treated with broad-spectrum antibiotics (Penicillin 100 U/L, ampicillin 1 g/L, neomycin 1 g/L, gentamycin 500 mg/L, metronidazole (0.5 mg/mL) and 1 mg/mL vancomycin) for two weeks prior to 12 Gy whole-body irradiation injury (12 Gy), and until harvesting intestines 3 days after injury. Mice were treated with EdU 1.5 hours prior to tissue collection.

(B) Staining for EdU showed less proliferation in DKO mice after injury compared to control. Hyperproliferative crypts (crypts with more than 5 $EdU^+$ cells) were counted for 50-100 well-oriented jejunal crypts per mouse and presented as % hyperproliferative crypts per total number of crypts mean ± SD (*$p < 0.05$, Ctrl versus DKO by unpaired two-tailed Welch's t-test; n = 3-4 mice/group). Scale bars, 50 µm

(C) IF staining for the stem cell maker SLC12A2 (green) with DAPI (blue) in jejunal sections, showing fewer $SLC12A2^+$ cells in DKO crypts after injury. Scale bars, 50 µm

(D) IF staining for the CBC marker and Notch target OLFM4 (green) with DAPI (blue) in jejunal sections, showing fewer $OFLM4^+$ cells in DKO crypts after injury. Scale bars, 50 µm
